# Coursing hyenas and stalking lions: the potential for inter- and intraspecific interactions

**DOI:** 10.1101/2022.02.23.481665

**Authors:** Nancy A. Barker, Francois G. Joubert, Marthin Kasaona, Gabriel Shatumbu, Vincent Stowbunenko, Kathleen A. Alexander, Rob Slotow, Wayne M. Getz

## Abstract

Resource partitioning promotes coexistence among guild members, and carnivores reduce interference competition through behavioural mechanisms that promote spatio-temporal separation. We analyzed sympatric lion and spotted hyena movements and activity patterns to ascertain the mechanisms facilitating their coexistence within semi-arid and wetland ecosystems. We identified recurrent high-use (revisitation) and extended stay (duration) areas within home ranges, and correlated environmental variables with movement-derived measures of inter- and intraspecific interactions. Spatial overlaps among lions and hyenas occurred at edges of home ranges, around water-points, along pathways between patches of high-use areas, and expanded during the wet season. Lions shared more of their home ranges with spotted hyenas in arid ecosystems, but shared more of their ranges with conspecifics in mesic environments. Despite shared space use, we found evidence for subtle temporal differences in the nocturnal movement and activity patterns between the two predators, suggesting a fine localized-scale avoidance strategy. Revisitation frequency and duration within home ranges were influenced by interspecific interactions, subsequent to land cover categories and diel cycles. Intraspecific interactions were also important for lions and, for hyenas, moon illumination and ungulates attracted to former anthrax carcass sites in Etosha, with distance to water in Chobe/Linyanti. Recursion and duration according to locales of competitor probabilities were similar among female lions and both sexes of hyenas, but different for male lions. Our results suggest that lions and spotted hyenas mediate the potential for interference competition through subtle differences in temporal activity, fine-scale habitat use differentiation, and localized reactive-avoidance behaviours. These findings enhance our understanding of the potential effects of interspecific interactions among large carnivore space-use patterns within an apex predator system, and show adaptability across heterogeneous and homogeneous environments. Future conservation plans should emphasize the importance of inter- and intraspecific competition within large carnivore communities, particularly moderating such effects within increasingly fragmented landscapes.

## Introduction

Spatio-temporal partitioning helps stabilize multi-species communities in which more than one species use the same resource [1–3]. More specifically, species whose ranges overlap forage different types of food, feed at different temporal schedules [4–9], demonstrate habitat separation, exhibit nonsynchronous spatial overlap or temporal partitioning [10–15]. Environmental heterogeneity provides temporary refugia where the risk of competition and injury is reduced [14, 16]. In addition, when there is an abundance of surplus resources the amount of food attracts numerous competitors, such that the energy required to exclude them becomes costly, and, therefore, competition ceases [17]. Resource use varies widely among sympatric carnivores [11,15,18–24], including African carnivores [3,8,25–32], and is presumed to promote coexistence [33–36].

Lions (*Panthera leo*) and spotted hyenas (*Crocuta crocuta*) are mainly crepuscular and nocturnal predators that demonstrate at least an 80% overlap between their daily activity budgets [13]. Although both species are sometimes active during cool winter days [37–39], they do not appear to use temporal partitioning to avoid interference competition [40]. The population densities of both lions and spotted hyenas are primarily influenced by the abundance of prey, and are, thus, positively correlated in some areas [41, 42]. As increasing prey abundance leads to an increase in the densities of both predators, however, the potential for interference competition increases with the likelihood of interspecific encounters [8]. Nonetheless, it appears that spotted hyenas derive benefits from sharing areas with lions. Hyenas appropriated up to 100% of lion kills in the Ngorongoro Crater when adult male lions were absent [43]. In the Amboseli National Park, hyenas did not avoid lion sounds from audio-call stations, and sometimes even approached these stations in response to lion roars [44]. This type of behaviour likely persists because avoiding lions may cost hyenas missed scavenging opportunities, given the high degree of overlap in their diets [45].

The ecological dynamics between lions and spotted hyenas are thus complex, and coexistence may be occurring due to the spatiotemporal partitioning of resources at fine spatial and temporal scales. Avoidance of potential competitors may be possible through small differences in the temporal use of habitats and shared resources [46]. Competitor avoidance is a behavioural strategy that reduces the probability of encounters within the foraging range of potential deadly rivals, thus enhancing the survivorship and fitness of the individual [47, 48]. However, avoidance of competitors is likely to invoke costs, such as a reduction in activity, a reduction in foraging rate or efficiency, or an increase in the use of refugia due to the perceived risk of predation [49–51].

Lions and hyenas appear to reduce some of the competitive effects of a shared diet by hunting prey of different sizes or ages [4,37,52,53]. Lions are able to hunt larger prey than spotted hyenas [54], but large groups of hyenas have adapted to hunting migratory prey populations in the Serengeti with the use of a unique commuting system [55]. Despite the lack of evidence for definite temporal partitioning in the activity periods of lions and hyenas, they exhibit slight differences during periods of activity. In several sites within Southern Africa and Tanzania, spotted hyenas were active for one continuous period during the night, while lions’ were active for two or three periods [13], whereas in the Southern Rift Valley of Kenya, hyenas were active after sunset and from the middle of the night to sunrise, with lions active throughout the night from 22h00 to dawn and after sunrise [56]. Therefore the two predators may actually be avoiding each other by utilizing the same prey abundant areas but at different times. In addition, food competition among lions and hyenas may be alleviated during periods of high resource availability, such as in the case of ungulate carcasses during anthrax outbreaks [57]. Furthermore, both species employ differences in their hunting behaviour, with hyenas mainly hunting large groups of prey and selecting target animals from rushing herds [37], while lions mainly employ stalk-and-ambush tactics of small herds of prey [39]. Thus, lions have the advantage in closed habitats while hyenas are likely to benefit from open habitats due to their cursorial nature. Lions tend to select for habitats with tall grass or steep embankments that promote hunting success and increases the catchability of prey [58–60], while hyenas seemingly appear to be able to utilize any type of habitat as habitat generalists [37, 38].

Although many studies have focused on the factors underlying spatio-temporal patterns in species distributions and resource use, few studies have examined the relationship between predator movement responses to each other to explain spatial overlap patterns. To our knowledge, studies have yet to elucidate fine scale movement and behavioural patterns in sympatric lions and spotted hyenas occurring in large scale natural systems that share much of the same resources, as mechanisms facilitating coexistence between these two predator species. This study fills this gap, by analysing fine-scaled movement data obtained from two different ecosystems subject to seasonal influxes of resources, to discern the behavioural differences that allow lions and spotted hyenas to co-exist. In particular, co-existence was assessed at the arid and mesic extremes of their environment, and the spatiotemporal or behavioural differences in their space use and activity patterns examined.

Data were collected using GPS radiocollars and activity accelerometers outfitted on lions and spotted hyenas located in both a large fenced National Park and within free-ranging areas of two distinct ecosystems. These data were analysed with the following objectives in mind: (1) to ascertain the environmental, bioclimatic and social factors influencing the spatiotemporal patterns of lions and spotted hyenas, while accounting for individual variation related to the age, sex, and condition of individuals; and, (2) to determine whether these were primarily influenced by heterospecific or homospecific competitors (i.e., conspecifics). We proceeded by first assessing the differences among the range use and proportion of shared space use among lion-hyena dyads between the two species. We then compared and contrasted the movement characteristics and activity patterns of the two species, and analyzed these movement patterns at various distances to competitors and conspecifics. Finally, we evaluated the relative roles of environmental variables versus inter- and intraspecific interactions in determining lion and hyena spatial distributions and movements. To investigate whether lion and hyena space-use patterns signify avoidance competition, we identified areas across lion and spotted hyena ranges with locations of higher-than-expected revisitation rates or locales of long-duration visits, and assessed whether these shifted in response to the presence of, or proximity to competitors. We also related these patterns to the distribution of resources at a landscape scale, including anthrax endemic areas in the arid environment.

## Materials and methods

### Ethics statement

Relevant permits required to carry out the research were obtained from the Ministry of Environment and Tourism, Namibia (Research/Collecting Permits 1724/2012, 1834/2013, 1956/2014) and from the Department of Wildlife and National Parks, Botswana (Research Permit EWT 8/36/4 XXVIII (35)). All animal handling procedures were conducted with the ethical clearance of the Animal Research Ethics Committee of the University of KwaZulu-Natal, South Africa (009/13/Animal), and the Institutional Animal Care and Use Committee of University of California at Berkeley (IACUC Protocol #R217-0512B) and Virginia Tech (IACUC Protocol # 15-012). Namibian specimens were shipped to RSA and Germany under CITES permits for the Regulations of Threatened or Protected Species (Permit/Certificate No. 0045192 and 157940).

### Study area

In brief, the study area covered 19,200km^2^ within the protected areas of the Southern Africa region: the Etosha National Park, a semi-arid savanna in northern Namibia; the Chobe National Park, Linyanti Conservancy, and the NG32 concession of the Okavango Delta, which comprises the Kalahari floodplains of northern Botswana (Fig 1, see S1 Appendix for additional details on these sites). In the Etosha National Park, certain regions are subject to an influx of seasonal resources from annual anthrax outbreaks [61]. The bacterial pathogen, *Bacillus anthracis*, is endemic as a major disease of various game species [62], and provides a significant subsidy of ungulate carcasses to predators and scavengers [57]. The Chobe-Linyanti region, hereafter “Chobe”, experiences a seasonal influx of migratory ungulate prey during the dry season in which they congregate around the perennial river [63]. Situated within the southeastern floodplains of the Okavango Delta, the NG32 concession is subject to an unimodal, annual flood pulse characterized by high variability in interannual flooding [64], and typically comprises a higher prey abundance during the dry season [65]. All study sites, therefore, have a season pulse in prey availability, although through different mechanisms.

**Fig 1.**
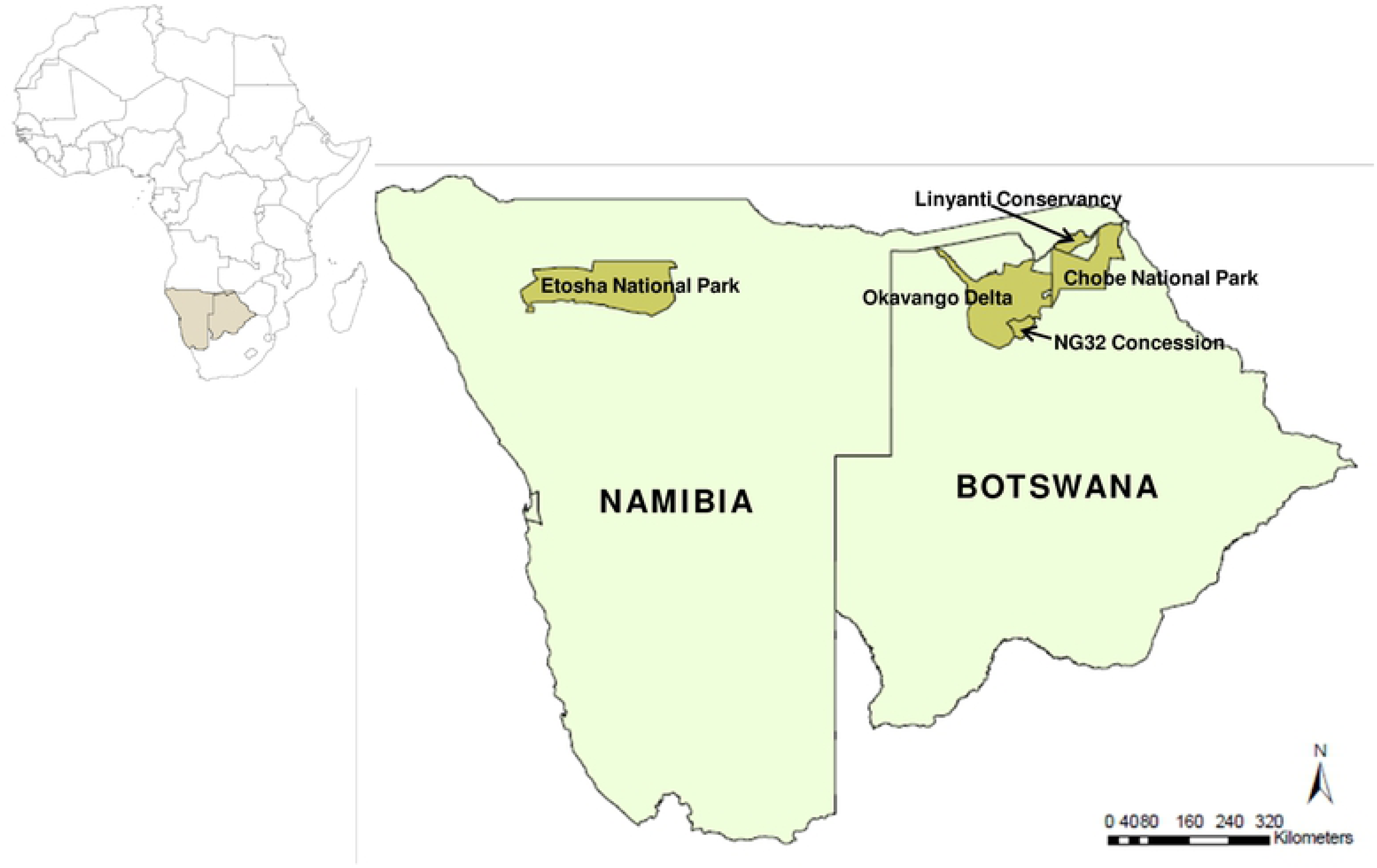
Location of the study areas. The map of the African continent shows the countries of Namibia and Botswana shaded, and the protected areas within these countries where the study was conducted. Maps were generated with ArcGIS (ESRI ArcMap v.10.0).

### Data collection

A total of 19 lions (13 females and 6 males) and 14 spotted hyenas (10 females and 4 males) were fitted with GPS-satellite collars with dual-axis accelerometers (IridiumTrackM, Lotek Wireless Inc., Newmarket, Ontario, Canada) (see S1 Appendix, Supporting information for additional details of collared animals and capture and sampling protocols). Collars were programmed to record GPS fixes on a schedule that consisted of a fix every 30 minutes during nocturnal periods (18h00 – 6h00 for Etosha individuals, and 17h00 – 8h00 for Botswana individuals), a fix every 5 minutes for two hours twice daily, once after sunset (19h00 – 21h00) and once before sunrise (4h00 – 6h00), and single diurnal fixes both at 10h00 and 14h00. For each datum, activity was averaged from acceleration collected in 8 second bursts over a duration of 240 seconds, and given a relative range between 0 and 255 (activity monitor values [AMVs]) to characterize the mean activity/acceleration. Relocation and activity data were downloaded from retrieved collars at the end of the study, with a subset of relocation data obtained from the satellite uplink of unretrieved collars via the Lotek web service (see S1 Appendix, Supporting information for details on data from unretrieved collars).

We retrieved 63% of all possible relocations while collars were deployed, due to the loss of collars from individuals that were killed, or for which we were unable to retrieve the collar (*n* = 3 lions and 6 hyenas). From the retrieved relocations (*n*=575,418) over all study sites, 73% were from lions with 27% from spotted hyenas. This dataset is roughly split between the two ecosystems with 43% of relocations from Etosha and 57% from Botswana. However, our relocation records from Etosha are significantly more complete (99.6% for lions and 84.1% for hyenas) compared to our relocation records from Botswana (53.3% for lions and 35.8% for hyenas). All statistical analyses were conducted in R version 3.5.1 (R Core Team, 2018), and all GIS applications were conducted in ArcGIS (ESRI ArcMap v.10.0, Redlands, CA, USA). See Table 1 for a list of expectations and key results.

**Table 1.**
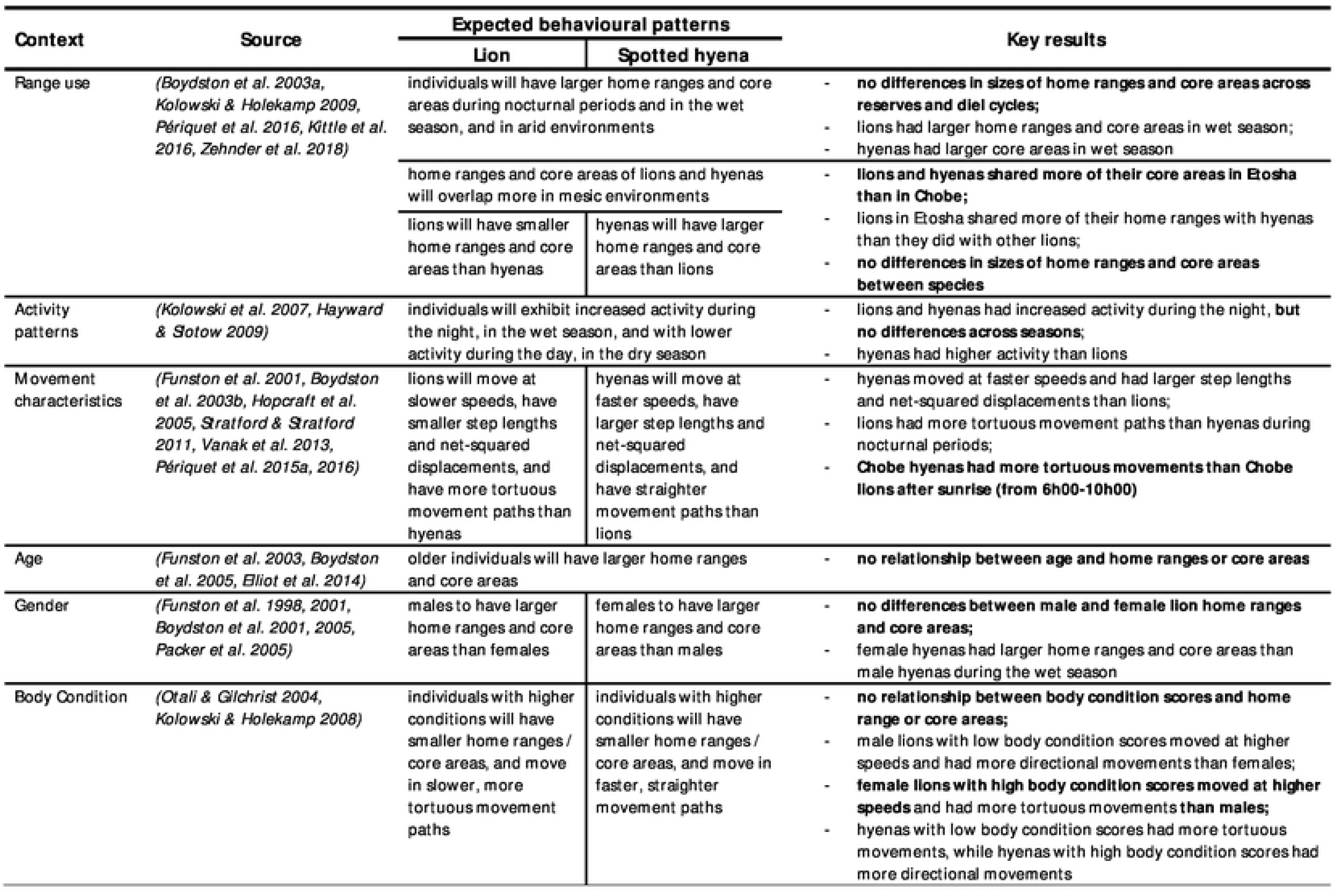
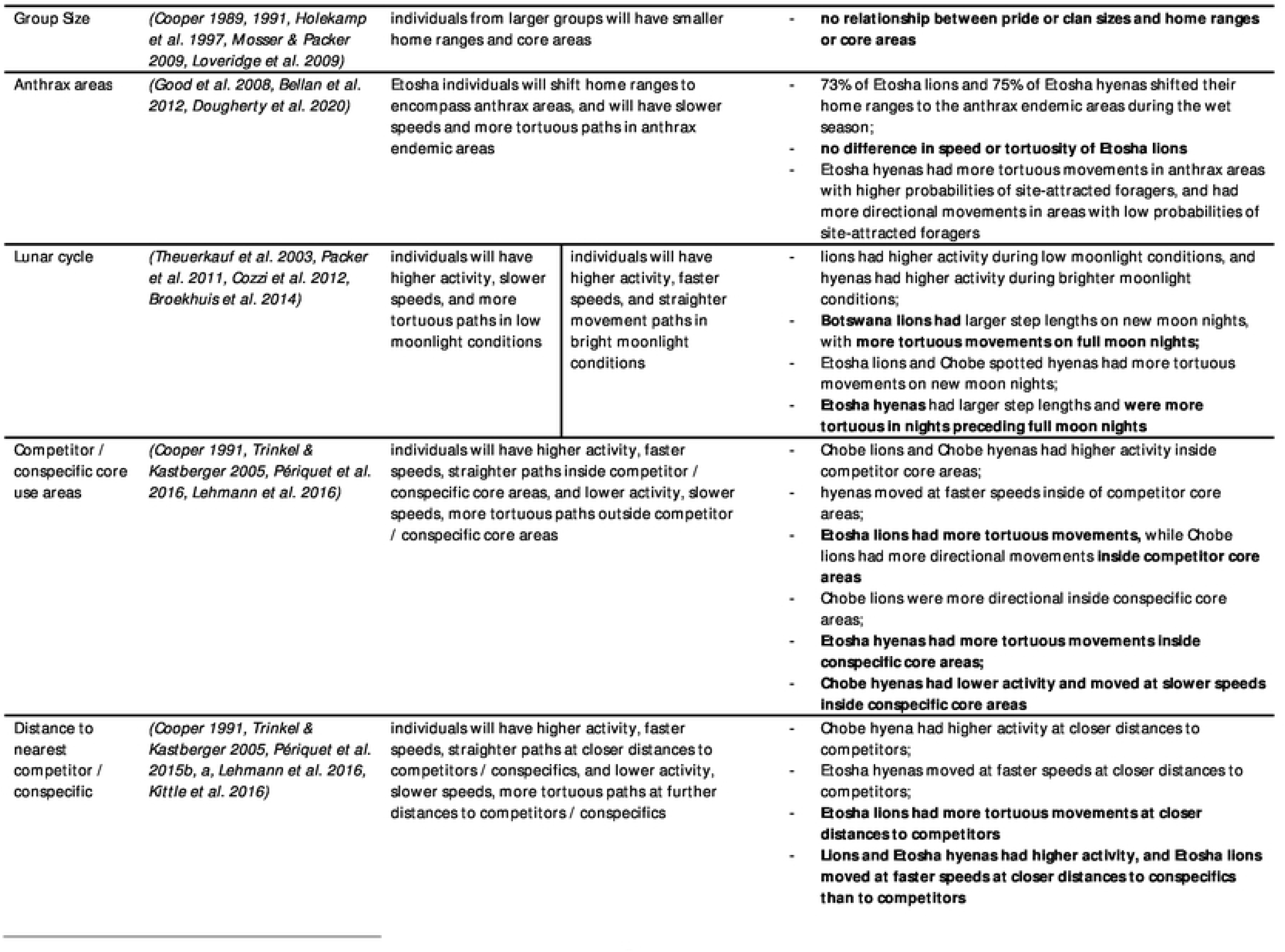
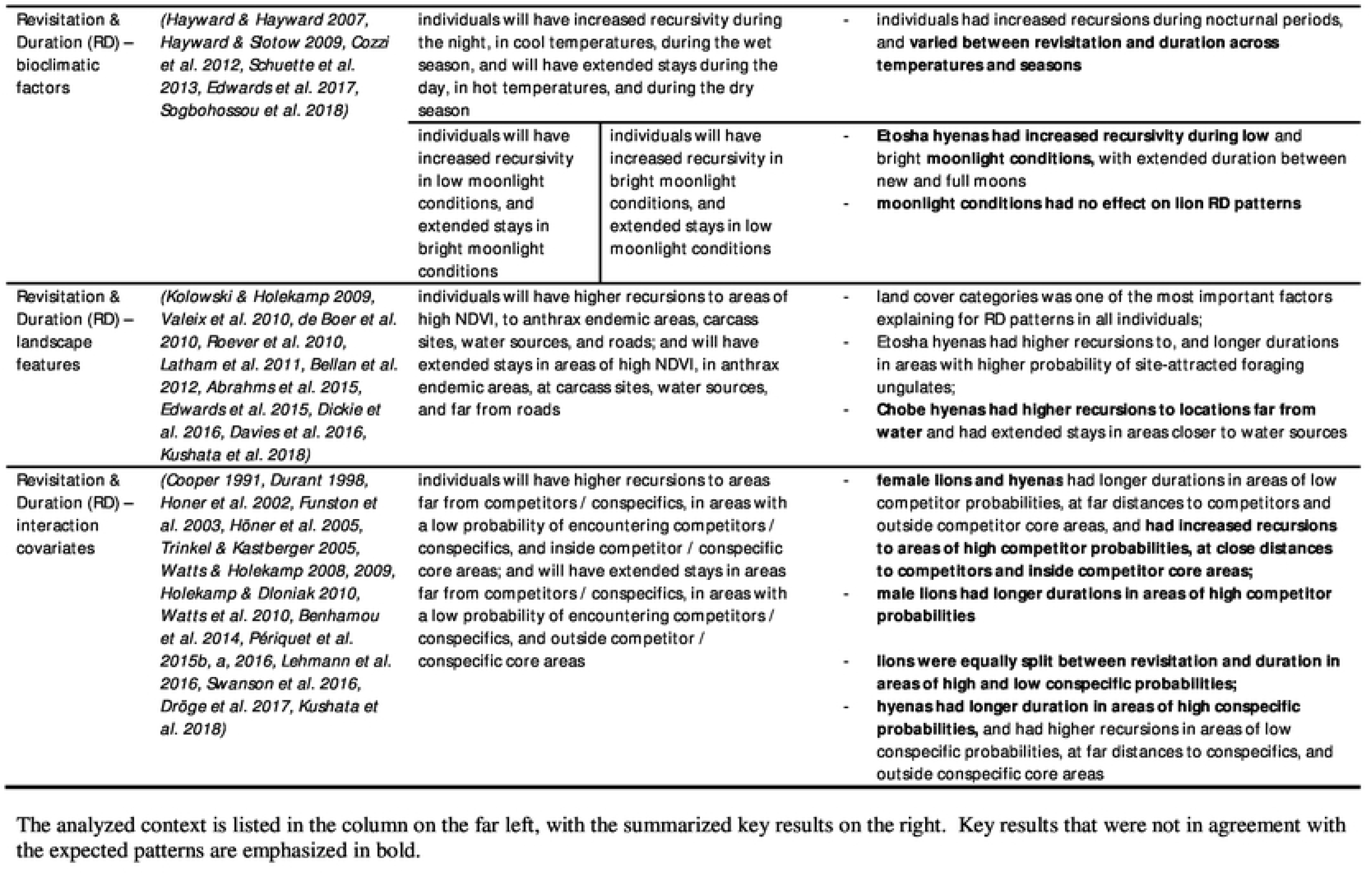
Expected behavioural patterns for lions and spotted hyenas, based on literature, for analyses undertaken in this study.

The analyzed context is listed in the column on the far left, with the summarized key results on the right. Key results that were not in agreement with the expected patterns are emphasized in bold.

### Species range use

The movement dataset of the lions and spotted hyenas were collected over a four year period split between the two study areas. From this dataset, we removed individuals with less than 30 tracking days, and created different subsets of sampling intervals to ensure for scale appropriateness in subsequent analysis [66]. Sampling intervals occurred every 30 or 5 minutes (depending on the sampling frequency over that sampling period, see above). The regularly sampled data downloaded from the collars all had some incidences of missed fixes (mean ± SE: lion 0.57 ± 0.03%, hyena 0.44 ± 0.06%), while the relocation fixes uploaded via satellites transmitted only a subset of locations during the study period in order to conserve collar batteries. Where collars were unable to be retrieved, movement data was downloaded from the Lotek web service, which consisted of relocations once every third fix of the programmed schedule (15 minute fixes during the 5 minute schedule, and 90 minute fixes during the 30 minute schedule). Thus the satellite data for some individuals (*n*=3 lions, *n*=6 hyenas) had missed fixes that ranged between 5 and 1040 minutes. Therefore, for analyses that included satellite individuals, we removed data that had a time difference in GPS fixes greater than 15 minutes from the 5 minute schedule, and greater than 90 minutes from the 30 minute schedule (which tended to occur towards the end of the study as a result of failing batteries). We then applied the “*ctmm*” R package [67] to fill in the missing coordinates in the schedule where required.

The data for analyses comprised of relocations from both collared individuals and relocations uploaded via satellite filled in with the ctmm method. From this data, nocturnal periods of relocations every 30 minutes were organized to include only fixes obtained during times of 18h00–6h00 or 17h00–8h00, and split into dry and wet seasons for the construction of home ranges and core areas. We filtered out 4-hour locations over a 24-hour period (maximum 6 fixes per day) from the relocation data, which were also split into dry and wet seasons to use for a comparison of nocturnal and diurnal ranges.

We constructed two types of utilization distributions (UDs) for each individual’s overall and seasonal ranges. First, we used the adehabitatHR kernel density estimator (KDE) with the reference bandwidth as the smoothing factor [68]. Second, we used the *a*-LoCoH (local convex hulls) adaptive method [69, 70], which facilitates the identification of regularly revisited sites, such as dens, waterholes, or riverbanks [69]. We used the 95% and 50% UDs (both methods) to represent respectively the home ranges and core use areas of individuals. We used the *t*-test in R v.3.5.1 to compare the sizes of home ranges and core areas among competitors (lions and spotted hyenas) with regards to location (Etosha versus Chobe), seasons, and segments of diel cycles. Within the kernel density ranges and utilization distributions, we used the intersection function from the “*rgeos*” R package [71] to compute the areas of overlap between neighbouring individual ranges, and used the *t*-test to determine the significance of differences in species’ overlapping ranges among ecosystems, across seasons and diel cycles.

### Species movement and activity patterns

To analyze the movement patterns of lions and spotted hyenas, we used the “*adehabitatLT*” R package [68] to calculate the step lengths and turning angles between successive locations. We calculated the speed and distance travelled (step length), path tortuosity (turning angle), and net-squared displacement (NSD) for each individual, and combined these individuals into species groups. Since turning angles range continuously on a circular scale from -180 to 180, we used the “*CircStats*” R package [72] to calculate the vectorized mean turning angle, which treats each observation as a vector on the unit circle to indicate the direction of the resultant vector [73]. Note that for CircStats, Watson’s two sample test is significant at *p* < 0.10. We then compared and contrasted the seasonal, sex-specific movement metrics of lions and spotted hyenas from each of the two ecosystems. We also assessed these differences over various times of the day, between diel cycles, land cover types, and across seasons.

We further analyzed the seasonal activity patterns of lions and spotted hyenas according to the full moon and new moon phases of the lunar cycle, as in Cozzi et al. [26]. We defined full moon nights when ≥95% of the lunar disc was illuminated and new moon nights were when moonlight intensity was ≤5%. For each day during full moon and new moon phases, we divided each 24-hour period into seven different sections (afternoon, dusk, night, nadir, night-end, dawn, morning) to reflect the main activity periods for lions and spotted hyenas [13]. We used “*suncalc*” in R [74] to calculate the times of periods for each day with dusk lasting from sundown to the end of evening astronomical twilight and dawn from the end of morning astronomical twilight to sunrise. The period between the end of evening twilight to the beginning of morning twilight was divided into three equal intervals in minutes to reflect night, nadir, and night-end. The day period from sunrise to sundown was divided from noon into morning and afternoon periods. We then calculated the proportion of the averaged activity measures that occurred during each of these periods.

For those individuals that had relocations within the core use area (or 50% isopleth) of either competitors or conspecifics, we assessed whether such proximity had any effect on the activity patterns as well as the speed and tortuosity of the focal animal. We assigned a value of 1 to each individual’s location that occurred within the core use area of a competitor’s or conspecific’s UD, and assigned a value of 0 to each individual’s location not within a competitor or conspecific core use area. We averaged the activity measures and movement metrics for each individual occurring inside and outside the core use areas of competitors and conspecifics. We then tested whether the average activity and movement metrics of lions and spotted hyenas differed significantly from when they were inside and outside of the core use areas of competitors and conspecifics.

### Inter- and intra-specific effects

#### Frequency of time-matched distances

The distances between collared individuals were obtained to analyze for interactive effects. We developed a temporally aligned matrix for each individual that overlapped in their collaring periods. For each sampling record of collar overlap, we measured the minimum Euclidean distance of that individual to all other collared individuals, at all locations, over the same time period. For each individual, we calculated the percentage frequency occurrences of time-matched distances between the individual to any heterospecific or homospecific competitor, in five frequency bins between distances of 5 km, 1 km, 200 m, 100 m, 50 m, and 10 m. We determined whether lions or spotted hyenas occurred at closer distances more often to each other (heterospecific competitors) than with one another among their own species (homospecific competitors).

#### Consecutive time points

In addition, we ascertained the length of time individuals spent with either competitors or conspecifics for all dyads that occurred at distances ≤ 5 km. We used the *distm* function from the “*geosphere*” R package to calculate the distance for each time-matched point in the trajectory of successive fixes for each dyad. We then calculated the number of consecutive time points (indicating a longer duration together) for which individuals of the dyad were at a distance below a given value (2 km, 1 km, 500 m, 200 m, 100 m). We again determined whether either lions or spotted hyenas occurred at closer distances more often to heterospecific or homospecific competitors, during consecutive time points.

#### Interaction variables

We included interaction variables to examine the influence of competitors and conspecifics on the rates of revisitations and visit durations of lions and spotted hyenas within their ranges. We measured distances between collared individuals and constructed GIS layers representing areas with a probability of competitor and conspecific use to analyze for interactive effects. To obtain the distances between collared individuals, we developed a temporally aligned matrix of each individual that overlapped with each other during the collaring period. For each sampling record of collar overlap, we measured the minimum Euclidean distance of that individual to all other collared individuals at all locations over the same times.

Individual seasonal UDs were overlaid and pixel cell values averaged to generate a total combined lion UD and a total combined spotted hyena UD for each of the dry and wet seasons. We converted the combined kernel UDs to volume UDs to obtain probability of use values for each cell. We subtracted volume UD values from 100 as in Kittle et al. [75], to obtain a more intuitive value with low use cells reflected by low values and high use cells reflected by high values. UD pixel values were then extracted and assigned to each locational point as a probability of competitor use area in ArcGIS v.10.0. We also repeated the process as above for a probability of conspecific use area. To construct the layer of the potential conspecific range, we overlaid the seasonal UDs of all other individuals of the same species, while excluding the individual the layer was being created for.

The Etosha lion UD was constructed from 11 individuals of 10 prides with 52,547 relocations for the dry season and 65,428 relocations for the wet season. The Chobe lion UD was constructed from 6 individuals of 5 prides with 30,016 relocations for the dry season and 43,356 relocations for the wet season. Etosha spotted hyena dry season UD was constructed from 57,211 relocations of 8 individuals from 8 clans, with wet season UD from 70,724 relocations of 7 individuals from 7 clans. Chobe spotted hyena dry season UD was constructed from 14,546 relocations of 4 individuals from 4 clans, with wet season UD from 15,335 relocations of 5 individuals from 5 clans.

### Ecogeographical variables

Ecogeographical variables (EGVs) such as distance to water, land cover types, and precipitation that have been statistically associated with species distribution [76–78], and were attached to each point within the RD space to evaluate their influences on lion and spotted hyena revisitations and visit durations (Table 2). We used ArcGIS (ESRI ArcMap v.10.0, Redlands, CA, USA) to measure the minimum Euclidean distance from all location points to various geographical features and landscape attributes, including distance to carcasses in Etosha (see S1 Appendix, Supporting information for additional details on the collection of Etosha carcass data).

**Table 2.** Description of ecogeographical variables (EGVs) used for cluster analyses.

| Ecogeographical Variables | Description | Data Resolution/Type |
| --- | --- | --- |
| *Bioclimatic variables* | Time of day | each relocation |
|  | Temperature | each relocation |
|  | Season | binary variable |
|  | Moon illumination | probability index, 0-1 |
|  | Precipitation amounts | pentad average |
| *Landscape features* | Slope | 30m x 30m |
|  | Land cover categories | 30m x 30m |
|  | NDVI | 30m x 30m, seasonal mean |
|  | Anthrax carcass probability | UD value |
|  | Distance to nearest available carcass | Euclidean, in m |
|  | Distance to nearest permanent water source | Euclidean, in m |
|  | Distance to nearest seasonal water source | Euclidean, in m |
|  | Distance to nearest road | Euclidean, in m |
| *Human disturbances* | Distance to nearest anthropogenic feature | Euclidean, in m |
| *Interspecific interactions\** | Distance to nearest competitor | Euclidean, in m |
|  | Competitor probability | UD value |
|  | Competitor core | 1 = inside core, 0 = outside core |
| *Intraspecific interactions\** | Distance to nearest conspecific | Euclidean, in m |
|  | Conspecific probability | UD value |
|  | Conspecific core | 1 = inside core, 0 = outside core |

EGVs were included in cluster analyses of the revisitation and duration (RD) space for each lion and spotted hyena individual within Etosha National Park, Namibia and the Chobe National Park, Linyanti Conservancy, and Okavango Delta, Botswana.
*\* not included in cluster analyses of lions from Okavango Delta, Botswana.*

Digital elevation maps of the four study areas were obtained from Landsat 8 images, courtesy of the U.S. Geological Survey, using a spatial resolution of 30 m. We derived the slope and aspect from these digital elevation maps in ArcGIS. Open-source land cover maps generated from LandSat thematic mapper data were downloaded for Namibia and Botswana (2010 Scheme II) via the RCMRD GeoPortal (http://geoportal.rcmrd.org), to which we assigned arbitrary values to reflect discrete land cover categories. We used Google Earth Engine’s [79] Normalized Difference Vegetation Index (NDVI) 8 day composites from the duration of the study period to obtain the mean NDVI values for each of the four study areas for each of the dry and wet season. As a measure of vegetation productivity [80–83], NDVI has been used as an index of prey availability [78, 84], with areas of increased vegetative cover correlated to increased ungulate and herbivore biomass [85–89]. These Landsat 8 composites are generated from Tier 1 orthorectified scenes, using the computed top-of-atmosphere reflectance. All the images from each 8-day period are included in the composite, with the most recent pixel as the composite value. NDVI values ranged from -1 to 1, with negative values corresponding to clouds and water, values near zero representing rock and bare soil, moderate values of 0.2 – 0.3 representing shrub and grassland, with high values close to 1 indicating temperate forests and tropical rainforests. Using ArcGIS, we calculated the mean center for each individual’s range, from which we obtained the relevant sunrise/set and moonrise/set times (Astronomical Applications Department of the U.S. Naval Observatory, from https://aa.usno.navy.mil), and interpolated the average precipitation from available CMAP Precipitation data (Climate Data and Resources, NOAA/OAR/ESRL PSD, Boulder, Colorado, USA, from https://www.esrl.noaa.gov.psd/). The “lunar” package in R version 3.5.1 (R Core Team, 2018) was used to assign moon phases and moon illumination values (ranging from 0 = new moon to 1 = full moon) to all locations, according to each location’s unique timestamp. For any location points collected between moonset and moonrise, the moon illumination was assigned a false logical vector.

### Time-use metrics

To assess how lions and spotted hyenas adjust their movements over time and across seasons, we quantified the time-use metrics (i.e., revisitation rates and visit duration) from the T-LoCoH hull parent points for each individual’s seasonal trajectories. We used the “*T-locoh.dev*” R package [70] to identify sites of repeated visits (nsv, number of separate visits to each cell) and sites of average visit durations (mnlv, mean number of locations per visit to each cell). A lion and spotted hyena whose ranges overlapped in Etosha is shown in Fig 2a, with all other individuals shown in S4 Fig (Supporting information). As we were interested in the period of time which both lions and spotted hyenas overlap in their activity periods [13], we chose to use an inter-visit gap (IVG) of 12 hours (one nocturnal period) to distinguish locations with more than 12 hours of time between them as separate visits. From this, we created density plots of each individual’s time use metrics on the landscape, which we refer to as revisitation and duration (RD) space (Fig 2b). We then obtained the local values of various ecogeographical and interaction variables (see below) and associated these covariate vectors to each point within the RD space.

**Fig 2.**
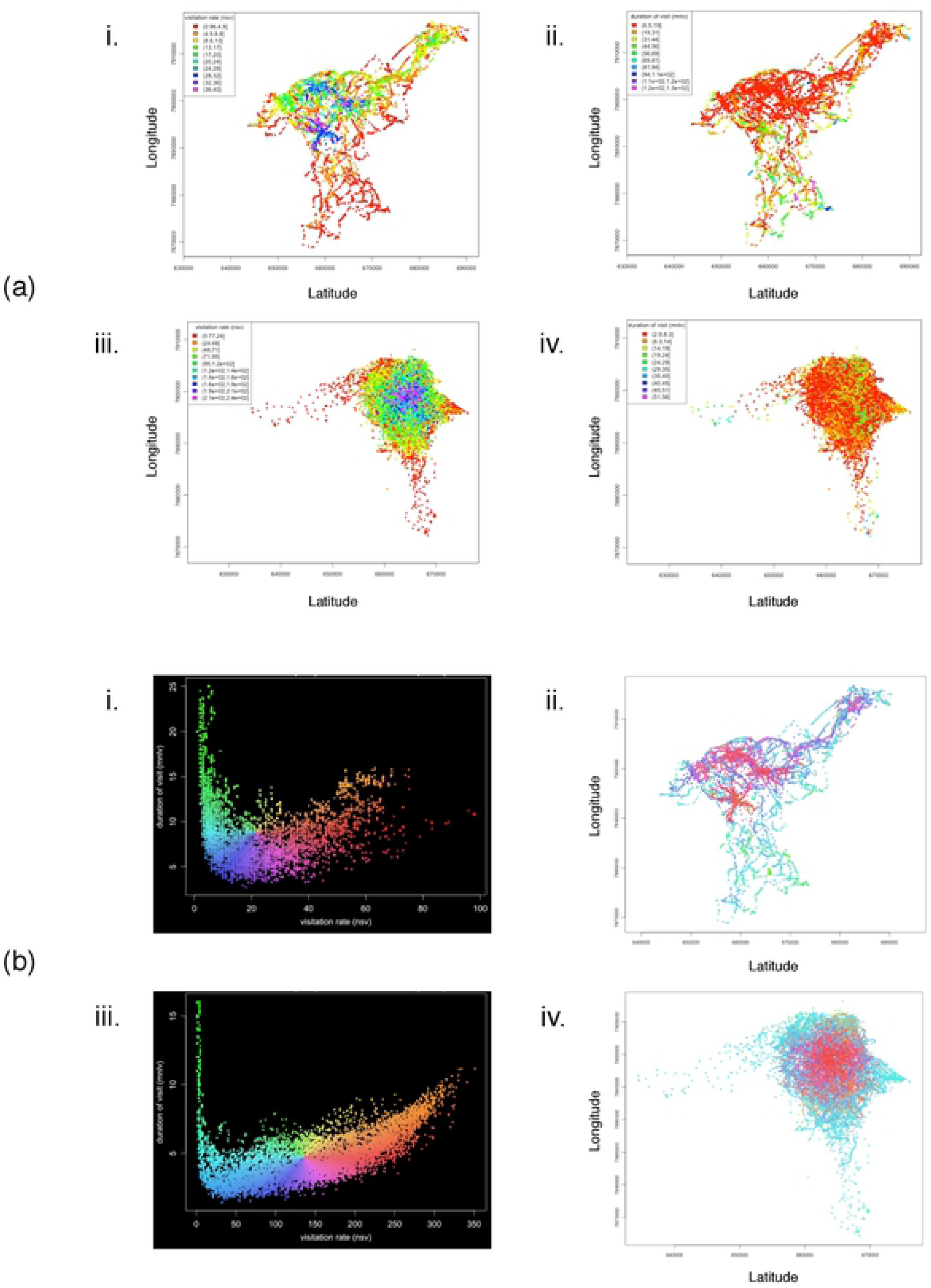
Revisitation and duration (RD) space plots indicated from the hull parent points of a lion and spotted hyena whose ranges overlapped. (a) Hull parent points for a collared female lion (NU-33865, i & ii, both figures) and female spotted hyena (GO-33869, iii & iv, both figures) whose ranges overlapped in Etosha. Parent points are coloured by visitation rate (nsv, number of separate visits; i & iii), and duration of visit (mnlv, mean number of locations in the hull per visit; ii & iv). (b) RD space scatterplots (i & iii) with X-axis = visitation rate (nsv), and Y-axis = duration of visit (mnlv), provide a legend for revisitation/duration (RD) values for the maps (ii & iv). Points in the RD space have been jiggled to better see point density, and each point within the RD space represents a hull (i) lion *n*=9160, (iii) hyena *n*=11898. Points on the maps (ii & iv) are coloured by their location in the RD space. Separate visits defined by an inter-visit gap period ≥ 12 hours. Hulls were created using the adaptive method. Duplicate points are offset by 1 map unit.

We used a factor analysis of mixed data (FAMD) to test for a statistically significant relationship between our selected ecogeographical and interaction variables with points in our constructed RD space. From the principal dimensions identified by our FAMD that describe >80% of the cumulative variation, we chose the variables with the highest scores in each dimension as the most important covariate. From this analysis, we determined the appropriate covariate combinations prior to clustering. We then performed a cluster analysis on the points within the RD space for each individual. We built mixed data type cluster models according to the FAMD results, and applied three different clustering algorithms for 2-8 clusters. We used a *k*-prototypes clustering algorithm [90] from the “*clustMixType*” R package, which is based on *k*-means for mixed type data. Due to the stochasticity of the *k*-prototype algorithm, each model configuration was recomputed 50 times with random initializations to obtain a model of minimum total distance. We also calculated a dissimilarity matrix using the “gower” metric in the daisy function from the “*cluster*” R package, which we used in the agglomerative hierarchical and PAM (partitioning around medoids) [91] clustering algorithms for further clustering. We colour-coded the points on the map according to the clustered results, and examined whether the points within a range of revisitation (R) and duration (D) values from the RD space fell mainly within the identified clusters. We then visually inspected the results of the clustering analysis and determined the appropriate clustering method. We chose the clustering method according to the distribution of the FAMD selected covariates for each cluster and the percentage of categories occurring in each cluster. We chose the *k*-prototype clustering algorithm as it resulted in more defined clustered groups within the individual’s RD space, and was more distinctive in the distribution of the clusters according to the covariates. An example is shown in Fig 3 for a lion and spotted hyena whose ranges overlapped in Etosha, with all other individuals shown in S5 Fig (Supporting information). We used the multivariate *t*-distribution algorithm from the “*ggplot*” R package to draw ellipses around the clustered points, and subsequently chose the number of clusters according to how the points were clustered together in the RD space.

**Fig 3.**
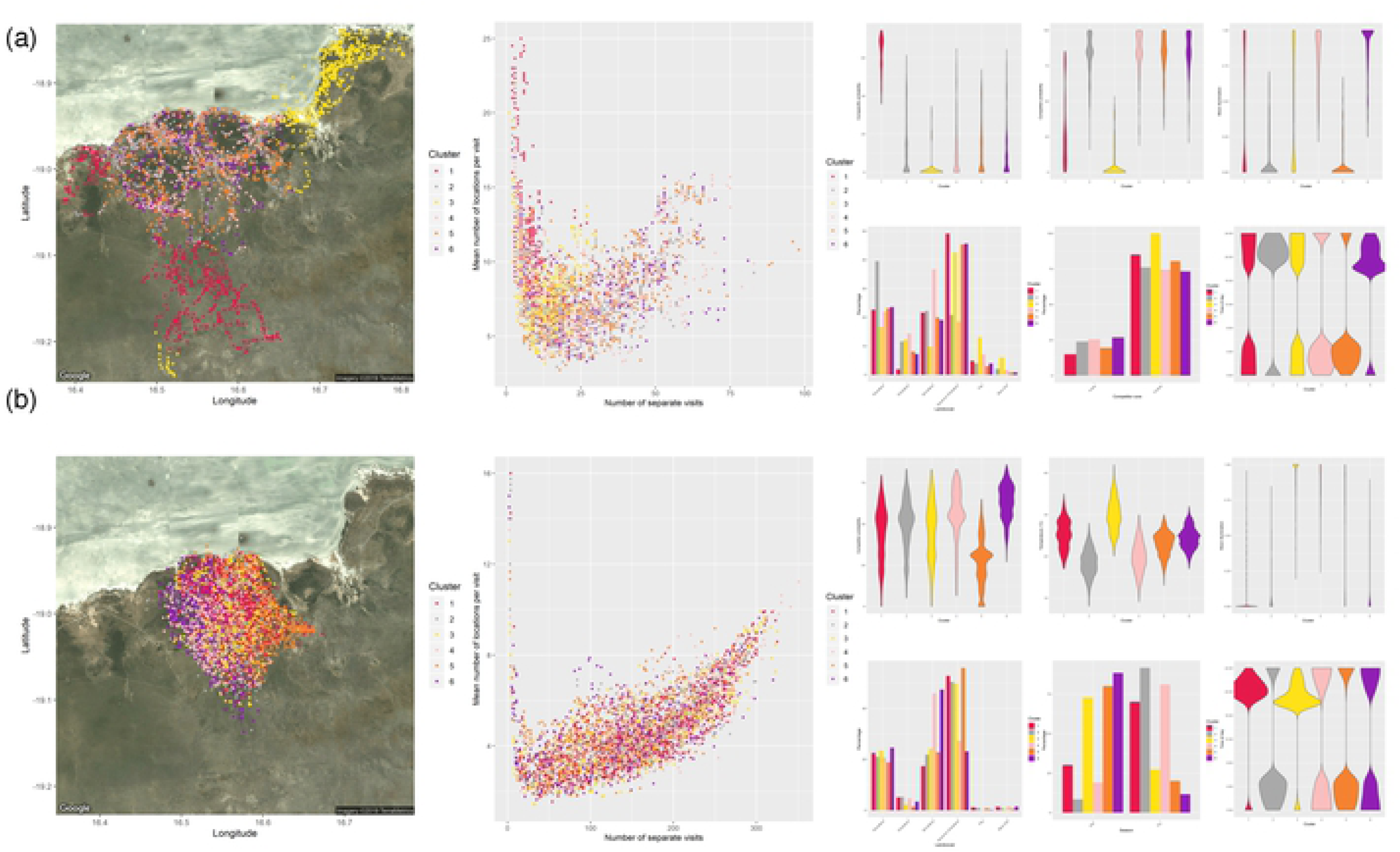
Cluster analyses of the revisitation and duration of a lion and spotted hyena whose ranges overlapped. Maps (far left panels) depict the individual relocations of a collared (a) lion (NU-33865) and (b) spotted hyena (GO-33869) whose ranges overlapped in Etosha. Relocations are colour-coded according to the clusters indicated by the range of revisitation (number of separate visits) and duration (mean number of locations per visit) values in RD space plots (central panels). Clusters in the RD space were determined with the *k*-prototype algorithm and are based on ecogeographical variables attached to each relocation. The smaller plots (right panels) present the distribution and percent category of clusters for each of the ecogeographical variables selected from the factor analysis of mixed data (FAMD).

## Results

### Species range use

Lions and spotted hyenas used the same types of habitat and occurred most frequently within similar land cover types between the two ecosystems. In Etosha, both species demonstrated higher frequencies of relocations within grassland habitats, whereas in Botswana sites, the most prevalent land cover types utilized by both species were shrublands (S1 Table, Supporting information). Kernel density UDs of lion and spotted hyena home ranges exhibited a high degree of spatial overlap within both Etosha and Chobe (Fig 4). Seasonal UDs for all individuals are presented in S2 Fig and S2 Table (Supporting information).

**Fig 4.**
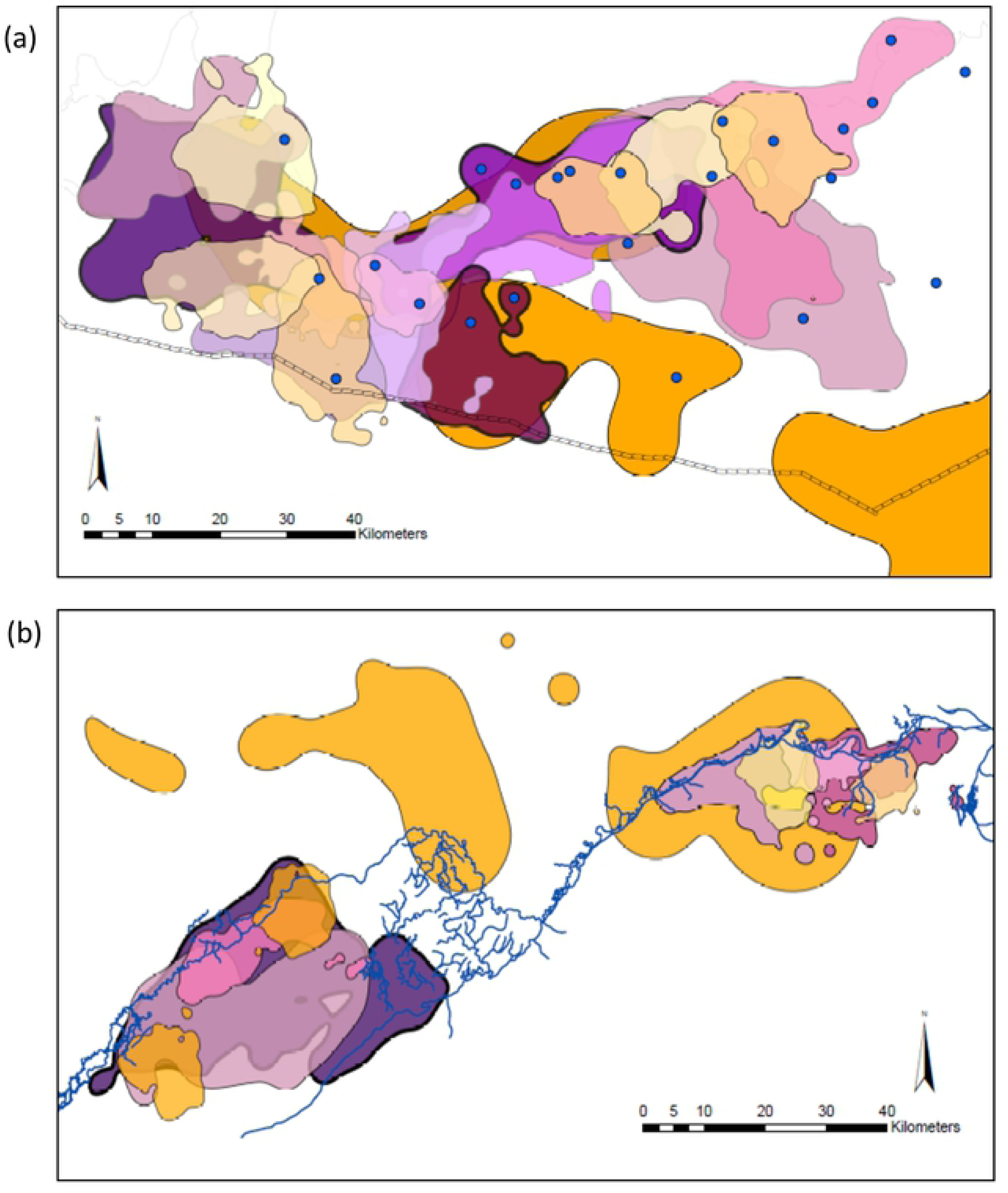
Overlapping ranges within study areas as represented by 95% kernel contours. Polygons of dark purple shades with black outlines = male lion ranges. Polygons of lighter purple/pink shades with grey outlines = female lion ranges. Polygons of orange/yellow shades = spotted hyena ranges. Overlapping polygons are set to 20% opacity for easier visualization. (a) Etosha site with the pan to the north and the park’s fence along the southern boundary. Blue dots represent permanent water points. (b) Chobe/Linyanti site with the river separating Botswana to the south and Namibia to the north. The river is indicated by blue lines.

In all cases, kernel density estimates of lion and spotted hyena home ranges were larger than the *a*-LoCoH estimates. Lion home ranges in Etosha estimated with the kernel method were (mean ± SE) 577.17 ± 93.90 km^2^, versus the *a*-LoCoH estimate of 361.96 ± 48.36 km^2^. In Botswana, lion home ranges estimated with the kernel method were 363.33 ± 119.81 km^2^, versus the *a*-LoCoH estimate of 257.31 ± 101.13 km^2^. Similarly, spotted hyena home ranges estimated with the kernel method were 718.56 ± 327.89 km^2^ in Etosha, and 478.68 ± 354.92 km^2^ in Chobe, while the *a*-LoCoH estimated hyena home ranges at 413.53 ± 115.07 km^2^ for Etosha and 194.08 ± 82.73 km^2^ for Chobe. Using the time-scaled distance measure incorporated into the UDs, there were no significant differences in the sizes of either *a*-LoCoH estimated 95% home ranges and 50% core use areas between lions and spotted hyenas during nocturnal and diurnal periods (mean ± SE lion: nocturnal home range 327.82 ± 49.61 km^2^, core area 98.98 ± 17.11 km^2^; diurnal home range 287.84 ± 49.81 km^2^, core area 95.64 ± 19.65 km^2^; hyena: nocturnal home range 331.49 ± 90.25 km^2^, core area 74.61 ± 15.84 km^2^; diurnal home range 257.47 ± 74.88 km^2^, core area 54.47 ± 12.08 km^2^; all *t*-tests, *p* > 0.05; S2 Table). There were also no significant differences in the sizes of either home ranges or core areas between lions and spotted hyenas in each reserve, regardless of seasons and circadian cycles (all *t*-tests, *p* > 0.05; S2 Table). However, when comparing conspecifics across reserves, Etosha lions had larger nocturnal core areas than Botswana lions (mean ± SE: Etosha 127.80 ± 24.42 km^2^, Botswana 62.95 ± 17.53 km^2^, *t* = -2.16, df = 15.4, *p* < 0.05). Similarly, Etosha hyenas had larger core areas than their counterparts in Chobe (mean ± SE: Etosha 85.21 ± 13.99 km^2^, Chobe 38.14 ± 21.56 km^2^, *t* = -2.77, df = 10.9, *p* < 0.05; S2 Table). Additionally, lions overall had larger home ranges and core areas in the wet season than they did in the dry season (mean ± SE diurnal home range: wet season 244.36 ± 42.33 km^2^, dry season 121.15 ± 29.72 km^2^, *t* = -2.38, df = 28.7, *p* < 0.05; nocturnal home range: wet season 278.39 ± 41.36 km^2^, dry season 162.45 ± 36.86 km^2^, *t* = -2.09, df = 30.8, *p* < 0.05; diurnal core area: wet season 95.69 ± 23.00 km^2^, dry season 42.23 ± 10.71 km^2^, *t* = -2.11, df = 22.6, *p* < 0.05; nocturnal core area: wet season 106.14 ± 21.40 km^2^, dry season 45.32 ± 11.65 km^2^, *t* = -2.50, df = 24.7, *p* < 0.05; S2 Table), whereas hyenas only had larger core areas in the wet season (mean ± SE: wet 90.54 ± 24.35 km^2^, dry 30.23 ± 6.93 km^2^, *t* = -2.38, df = 12.8, *p* < 0.05; S2 Table).

The total area of overlap in the home ranges and core areas of the two predators, both the other’s main competitor, are also presented for each pair in S3 Table (Supporting information). We calculated the proportion of lion home ranges and their core areas overlapped by spotted hyena home ranges and core areas, as well as the proportion of spotted hyena home ranges and their core areas overlapped by lion ranges and core areas (S4 Table, Supporting information). Lions in Etosha shared up to (mean ± SE) 20 ± 4% of their home ranges and 8 ± 2% of their core use areas with hyenas. Meanwhile, hyenas shared up to 17 ± 3% of their home range and 8 ± 2% of their core use areas with lions in Etosha. Similarly in Chobe, lions shared up to (mean ± SE) 23 ± 5% of their home ranges and 3 ± 1% of their core use areas with spotted hyenas, while hyenas shared up to 23 ± 4% of their home ranges, and less than 1 ± 0.5% of their core use areas with lions. The proportion of overlap in the core use areas between lions and hyenas were larger in Etosha than they were in Chobe (mean ± SE: Etosha 8 ± 2%, Chobe 3 ± 1%, *t* = 2.19, df = 81.0, *p* < 0.05; S4 Table). In addition, the proportion of overlap in the core use areas between lions and hyenas in Etosha were larger during the wet season than they were for the dry season (mean ± SE: wet season 14 ± 4%, dry season 3 ± 2%, *t* = -2.31, df = 34.7, *p* < 0.05; S4 Table). Although lions did not differ in the sizes of overlapped areas between conspecifics or competitors, a larger proportion of Etosha lion home ranges overlapped with competitors than they did with conspecifics (mean ± SE: competitors 20 ± 4%, conspecifics 10 ± 2%, *t* = -2.35, df = 103.7, *p* < 0.05; S4 Table). By contrast, lion conspecifics in Chobe shared significantly more of their home ranges than they did with conspecifics in Etosha (mean ± SE: Etosha 10 ± 2%, Chobe 26 ± 7%, *t* = -2.19, df = 28.9, *p* < 0.05; S4 Table).

### Species movement and activity patterns

Descriptive analyses of lion and spotted hyena movement descriptors are presented in S2 Appendix and S5 Table (Supporting information). Despite being temporally aligned with the activity periods of lions at night, spotted hyenas exhibited nearly twice the activity rates of lions (mean ± SE: lion 21.88 ± 10.3 AMVs, hyena 40.20 ± 24.0 AMVs, *t* = -2.90, df = 7.2, *p* < 0.05; Fig 5; S5 Table). They also moved at characteristically higher speeds than lions in both ecosystems (Etosha mean ± SE: lion 0.187 ± 0.08 m/s, hyena 0.364 ± 0.14 m/s; Chobe mean ± SE: lion 0.128 ± 0.08 m/s, hyena 0.310 ± 0.16 m/s; S2 Appendix), and had greater nocturnal mean step lengths (mean ± SE: lion 288.90 ± 99.84 m, hyena 618.12 ± 189.85 m; all *t*-tests, *p* < 0.001; S2 Appendix; S5 Table). Spotted hyenas were consistent in that they moved at significantly faster speeds and travelled significantly further than lions throughout different time periods and across both ecosystems (all *t*-tests, *p* < 0.05; S2 Appendix; S5 Table), except for during the wet season in Chobe. Although 62% of lions had greater step lengths during dawn than they did dusk (χ*^2^* = 5.33, df = 1, *p* < 0.05), lions travelled significantly further in dusk periods than throughout dawn periods (mean ± SE: at dusk 64.79 ± 2.29 m, at dawn 50.64 ± 3.87 m; *t* = 3.14, df = 9.7, *p* < 0.05; S2 Appendix), while hyenas did not show any differences between the two periods.

**Fig 5.**
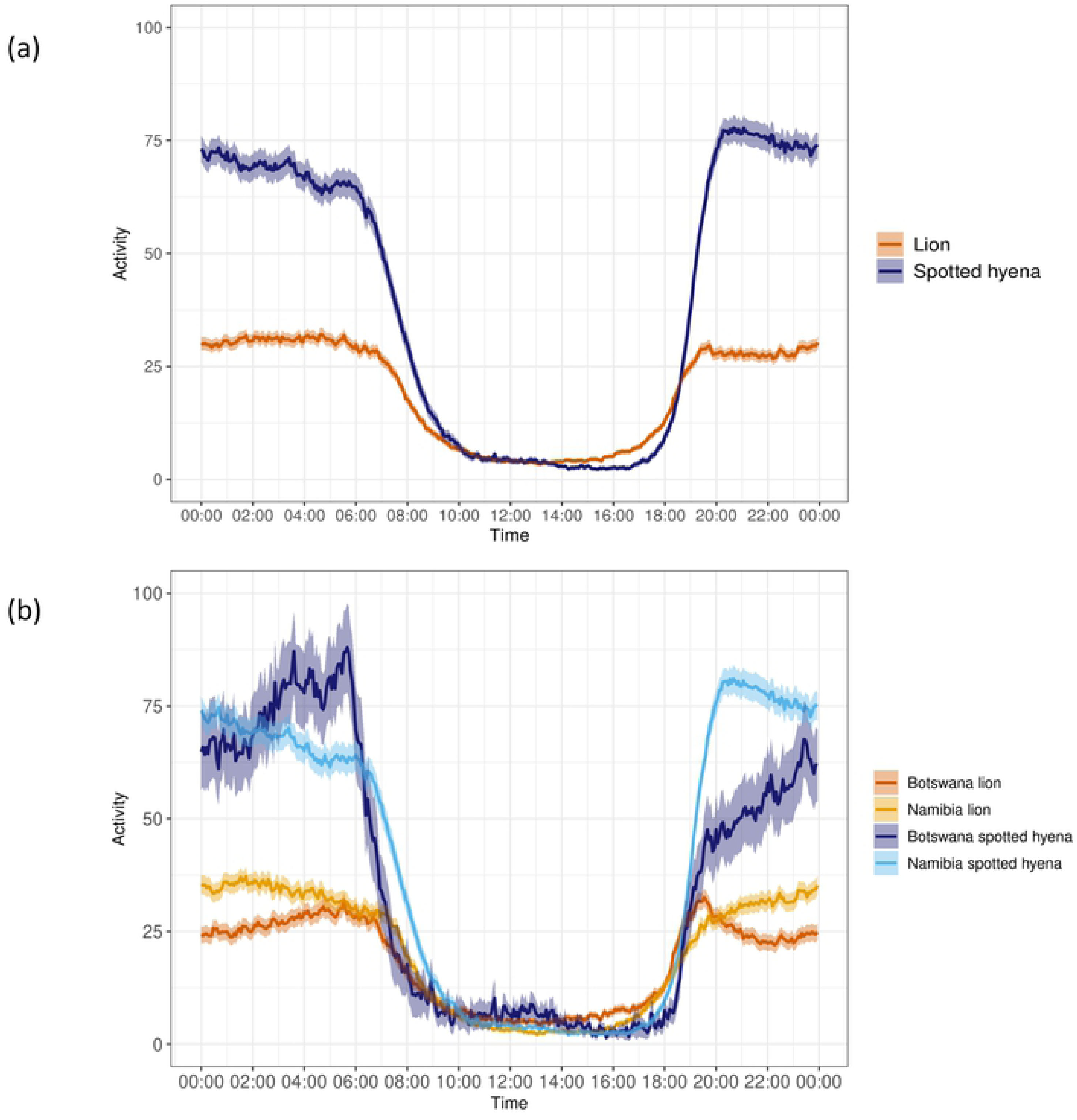
Mean activity of lions and spotted hyenas. Activity rates over the 24-hour cycle of (a) lions and spotted hyenas, and (b) lions and spotted hyenas from the Etosha National Park, Namibia and the Chobe National Park, Linyanti Conservancy, and the Okavango Delta^†^, Botswana. Means are represented by solid lines with 95% confidence intervals the shaded bars. ^†^*No spotted hyenas were collared from the Okavango Delta, Botswana*.

During 24-hour periods, spotted hyenas demonstrated more directional movements in the semi-arid Etosha ecosystem, and had more tortuous movements in the wetland environments of Chobe (dry season mean ± SE: Etosha -2.012 ± 0.89, Chobe 2.660 ± 1.01, Watson’s test statistic: 0.217, *p* < 0.05; wet season mean ± SE: Etosha 1.637 ± 0.94, Chobe -2.942 ± 0.90, Watson’s test statistic: 0.200, *p* < 0.05; S2 Appendix; S5 Table). Contrarily, hyenas exhibited more directional movements in Chobe and more tortuous movements in Etosha during dusk and dawn periods of the wet season (mean ± SE: Chobe 0.004 ± 0.56, Etosha 0.012 ± 0.58, Watson’s test statistic: 0.171, 0.05 < *p* < 0.10; S2 Appendix; S5 Table.) Similarly, lions had more directional movements in Etosha than they did in Botswana during the dry season (mean ± SE: Etosha 0.018 ± 0.85, Botswana -0.068 ± 0.86, Watson’s test statistic: 0.177, 0.05 < *p* < 0.10; S2 Appendix; S5 Table). In addition, lions were also more directional than hyenas over the 24-hour period in Etosha (mean ± SE: lion 0.080 ± 0.62, hyena -2.852 ± 0.82, Watson’s test statistic: 0.305, *p* < 0.01), and in Chobe during the wet season (mean ± SE: lion 0.209 ± 0.83, hyena -2.942 ± 0.90, Watson’s test statistic: 0.189, *p* < 0.05; S2 Appendix; S5 Table). However, lions were more tortuous than hyenas were in Etosha during nocturnal and dusk/dawn periods (mean ± SE: lion 0.053 ± 0.73, hyena 0.013 ± 0.56, Watson’s test statistic: 0.305, *p* < 0.01; S2 Appendix; S5 Table), while in Chobe, hyenas had more tortuous movements than lions during the late morning (Watson’s test statistic 0.269, *p* < 0.05; S2 Appendix 10).

Moreover, lion and hyena movements and activity were found to differ according to the lunar cycle. During nocturnal periods in Etosha, lions travelled further and had higher activity during low light conditions (i.e., waxing and waning crescents; step length *F* = 16.36, activity *F* = 11.33, *p* < 0.0001; S5 Table). In the dry season, lions in Etosha demonstrated more tortuous movements on new moon nights (mean ± SE: new moon 0.351 ± 0.86, full moon 0.013 ± 0.84, *F* = 22.99, *p* < 0.05; S5 Table), and had more directional movements during periods of low light conditions (waxing, waning crescents; Etosha lion: *F* = 34.10, *p* < 0.0001; S2 Appendix), in addition to full moon nights. However, hyena activity in Etosha demonstrated two peaks during waxing gibbous to full moon nights, and again during new moon nights, with decreased activity during first quarter phases and after full moon nights (*F* = 20.13, *p* < 0.0001; S5 Table). During the wet season in Etosha, hyenas had more directional movements during the brightest phases (i.e., waxing/waning gibbous and full moon), and exhibited more tortuous movements during new moon nights, and first and last quarter phases (*F* = 2.32, *p* < 0.05; S5 Table).

Similarly, lions in Botswana travelled further and had higher activity on new moon nights (mean ± SE: step length, new moon 242.81 ± 146.57 m, full moon 222.71 ± 139.56 m, *F* = 7.34; activity, new moon 29.34 ± 45.53, full moon 27.16 ± 44.46, *F* = 19.39, *p* < 0.05; S5 Table), although they had more tortuous movements during full moon nights (mean ± SE: full moon 0.442 ± 0.91, new moon -0.272 ± 0.80; Watson’s test statistic: 0.168 and 0.5 < *p* < 0.10; S5 Table). In addition, during dusk/dawn periods of the wet season, Botswana lions had mostly tortuous movements in first quarter and waning gibbous, with mostly directional movements in waxing gibbous phases (*F* = 2.51, *p* < 0.05; S5 Table). Contrarily, hyenas in Chobe had mostly directional movements during full moon nights and were more tortuous during new moon nights during nocturnal periods of the dry season (mean ± SE: full moon 0.020 ± 0.96, new moon 0.139 ± 0.94, *F* = 4.19, *p* < 0.05; S5 Table).

Furthermore, the proportion of activity as shown by lions and spotted hyenas according to the lunar cycle during nocturnal periods reveals a seasonal effect and regional differences on their temporal activity patterns. For both the arid and mesic environments, lions and spotted hyenas exhibited significantly higher proportions of activity during the periods of the night from dusk to dawn (Night mean ± SE: lion 0.531 ± 0.014 AMVs, hyena 0.591 ± 0.009 AMVs; Dusk/dawn mean ± SE: lion 0.356 ± 0.010 AMVs, hyena 0.350 ± 0.007 AMVs; *t* = -16.25, df = 23.1, *p* < 0.0001; Fig 6), regardless of moon phase.

**Fig 6.**
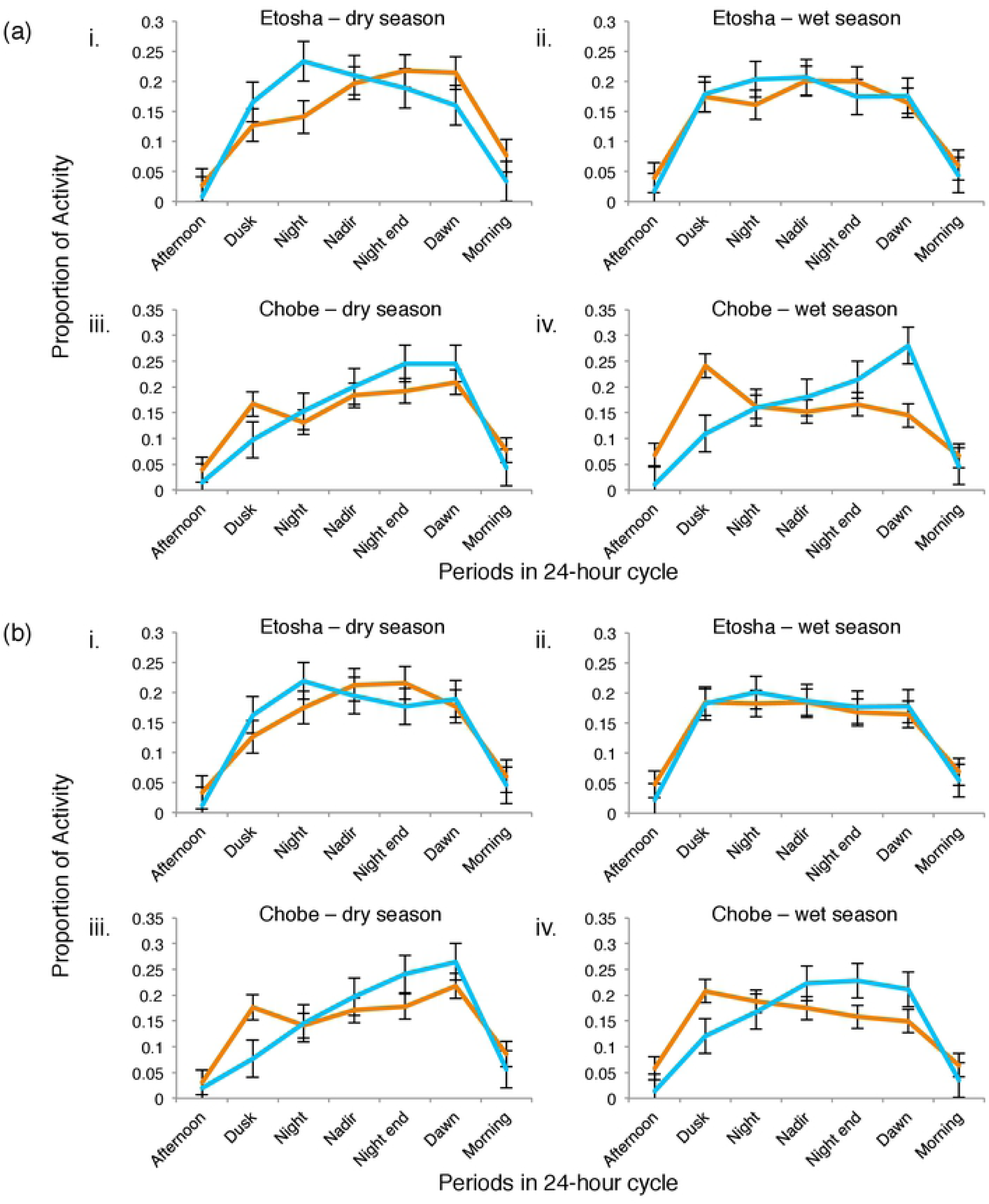
Seasonal activity of lions and spotted hyenas according to full and new moons. Proportion of activity of lions (orange lines) and spotted hyenas (blue lines) during full moon (a) and new moon (b) phases for each of the dry and wet seasons in Etosha and Chobe/Linyanti study areas. The 24hr cycle was subdivided into seven different periods. Night/nadir/night-end consists of the end of evening twilight to the beginning of morning twilight divided into three equal intervals. Afternoon = noon to sundown; dusk = sundown to twilight end; dawn = beginning of morning twilight to sunrise; morning = sunrise to noon. Points represent the mean and error bars the standard error (SE).

During both new and full moon nights in Etosha, spotted hyenas exhibit heightened proportions of activity in the initial phase of the night (dusk-night mean ± SE: lion 0.159 ± 0.008 AMVs, hyena 0.193 ± 0.009 AMVs, *t* = -2.77, df = 14, *p* < 0.05; Fig 6; S5 Table). However, the opposite is true for Chobe with lions having heightened proportions of activity in the initial phase (dusk-night mean ± SE: lion 0.177 ± 0.013 AMVs, hyena 0.129 ± 0.012 AMVs, *t* = -2.81, df = 13.9, *p* < 0.05; Fig 6; S5 Table), with hyenas having higher proportions of activity during the latter phase (night end-dawn mean ± SE: lion 0.177 ± 0.010 AMVs, hyena 0.241 ± 0.008 AMVs, *t* = -5.03, df = 13.7, *p* < 0.001; Fig 6; S5 Table). Despite whether lions or hyenas were exhibiting higher proportions of activity during certain periods of the night, there remains a temporal shift in activity between lions and hyenas in which periods of higher proportions of activity is dominated by one species during different time periods of the night. Thus, these distinctive differences in the activity patterns between the two species during the night suggest a temporal partitioning strategy in areas where both species co-exist.

### Inter- and intra-specific effects

#### Frequency of time-matched distances

Of 1,155,488 records of distances between collared individuals that overlapped in time, 46.7% occurred between conspecifics (26.1% for lions and 20.6% for spotted hyenas), and 53.3% occurred between lions and spotted hyenas (competitors). From these data, we extracted all measured records between two collared individuals that occurred at a distance of ≤5 km (*n* = 34,682). Overall, predators were at distances of ≤5 km to competitors and conspecifics with nearly equal frequency (51% and 49%; *x^2^* = 0.071, df = 1, *p* > 0.05; Fig 7). Etosha lions occurred at distances of ≤5 km with each other more often than Chobe lions did (*x^2^* = 39.69, df = 1, *p* < 0.001), while Chobe hyenas were at distances of ≤5 km with each other more often than Etosha hyenas were (*x^2^* = 8.29, df = 1, *p* < 0.005). Etosha lions occurred more often with competitors at further distances (1-5 km), and with conspecifics at closer distances (0-50 m), whereas hyenas occurred at distances of <5 km with competitors more often than with conspecifics (Fig 7, and S6 Table, Supporting information). However, hyenas in Chobe occurred more often with conspecifics than they did with competitors at distances of 200 m-1 km. In addition, lions and hyenas tended to be at further distances to competitors than to conspecifics, although this was not significant in all cases (S6 Table).

**Fig 7.**
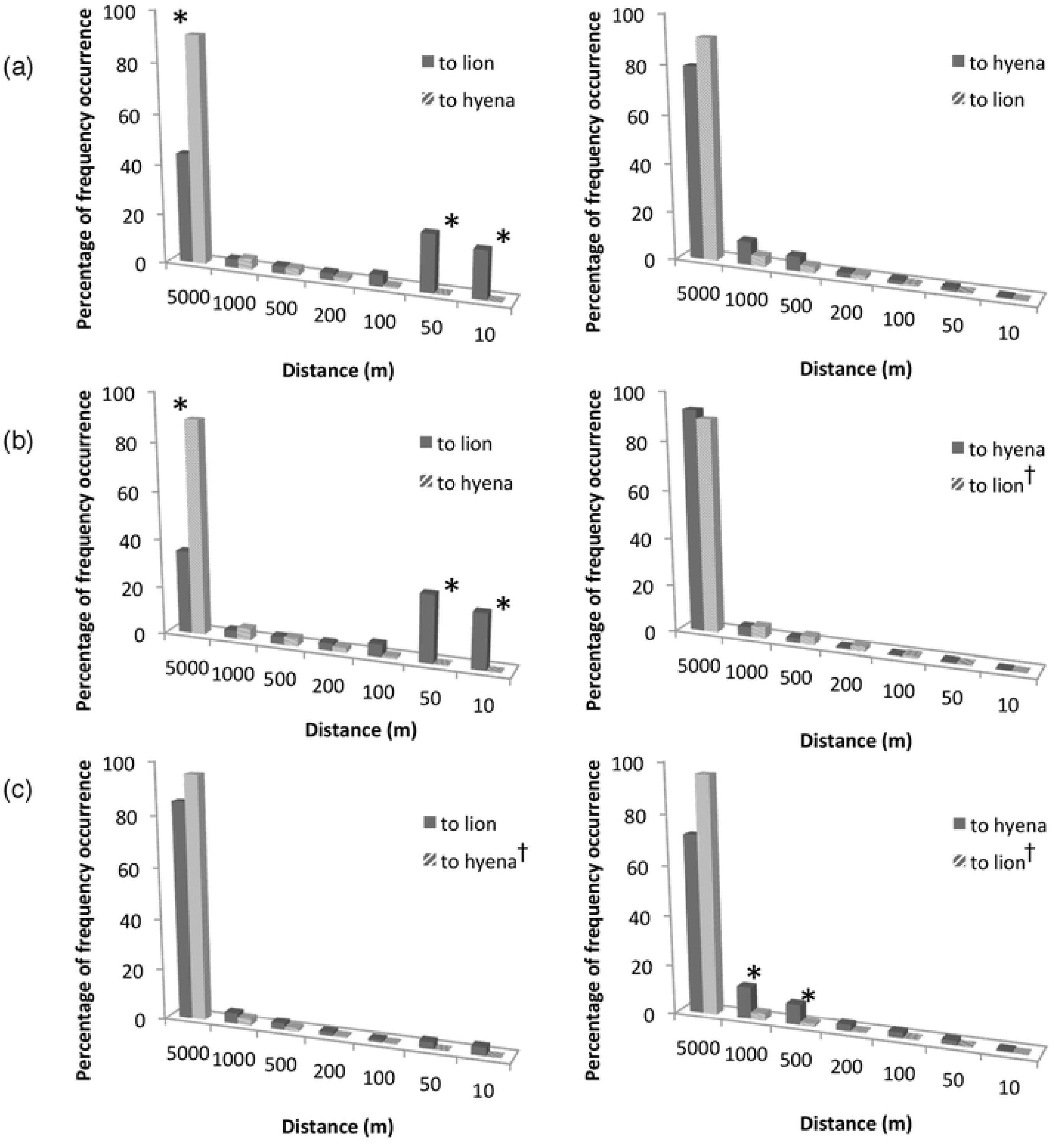
Frequency of time-matched distances between lions and spotted hyenas. Percent frequency occurrence of time-matched distances from lions (left side figures) and spotted hyenas (right side figures) to collared individuals at distance intervals of 0-10 m, >10-50 m, >50-100 m, >100-200 m, >200-500 m, >500-1000 m, and >1-5 km. Dark grey bars reflect frequency of distances to conspecifics, and hatched bars frequency of distances to competitors. (a) Overall percentage frequency occurrence, (b) Etosha groups, and (c) Chobe/Linyanti groups. Asterisks denotes the significant difference between competing groups for that interval; and a dagger indicates which competing group has the greater percent frequency occurrence between 0-5 km. Statistical results are presented in S6 Table (Supporting information).

#### Consecutive time points

During consecutive time points (which indicates a longer duration of time together), there were significantly more instances of lion-lion dyads at distance intervals of 0-100 m, 100-200 m, 200-500 m, and 500-1000 m than there were of lion-hyena dyads, indicating that lions spend more time together at these distances than they do with hyenas (Fig 8). In addition, intraspecific dyads have more instances (85%) of being together for consecutive time points than interspecific dyads (15%). Lion dyads consistently presented higher values for both the “11-30” and “>30” consecutive time point groups, indicating that they spend more time together at these distances than they did with hyenas, and more than hyenas did (S7 Table, Supporting information). However, despite higher frequencies of hyenas spending more time together than they did with competitors at shorter consecutive intervals (i.e. “1” and “2” groups indicating 5-10 mins), there were more occurrences of lion-hyena dyads at greater consecutive time points (“>30” group), indicating a longer time interval of at least 300 consecutive minutes for lion-hyena dyads at distances between 0-2 km.

**Fig 8.**
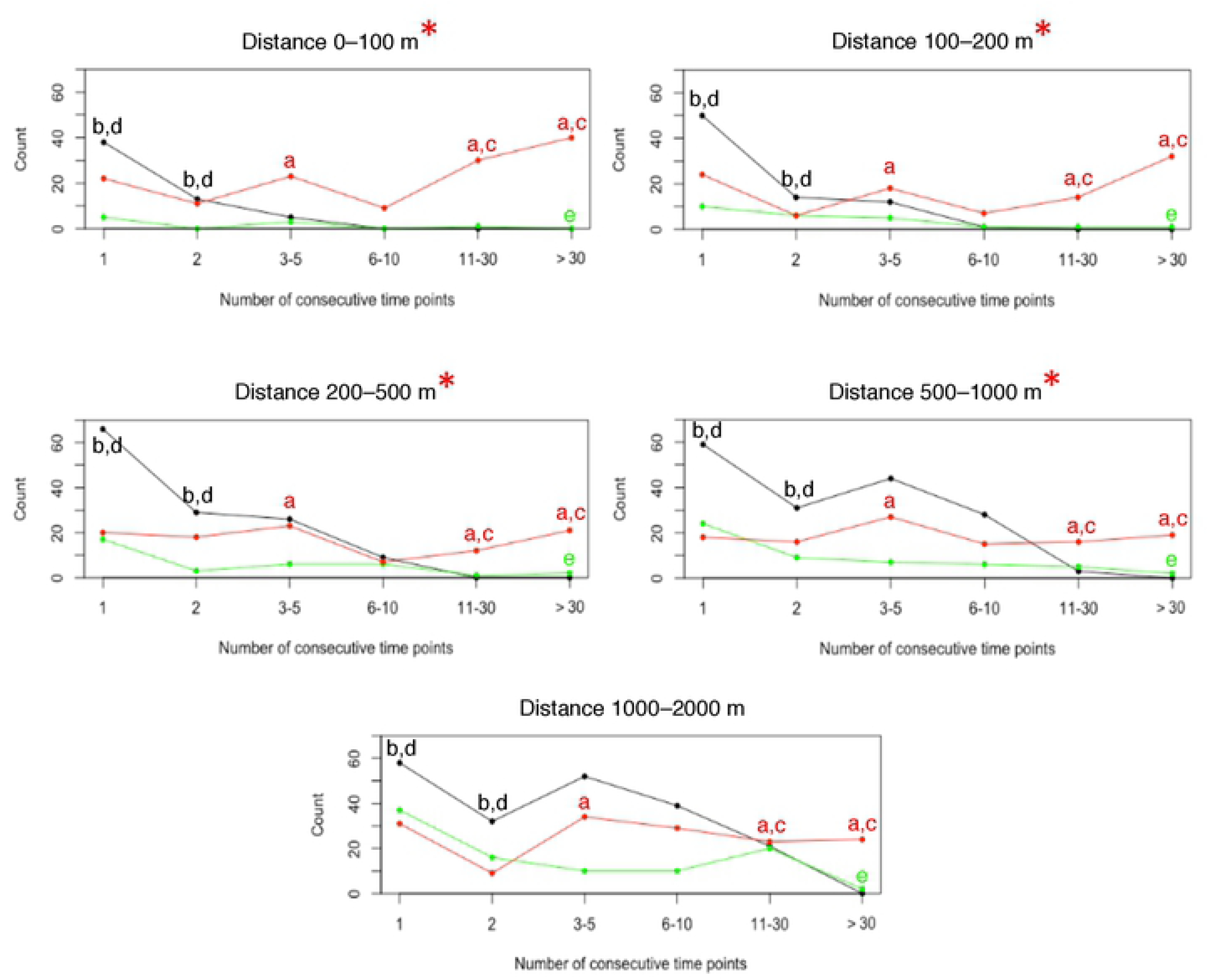
Consecutive time points between lions and spotted hyenas at different distance intervals. Number of consecutive time points (indicating longer time duration) for which pairs of lion-lion (red), hyena-hyena (black), and lion-hyena (green) dyads were at distances of 0-100, >100-200, >200-500, >500-1000, or >1000-2000 m. Red asterisks at distance intervals indicates where the lion-lion dyad had significantly more consecutive time points contrasted to lion-hyena dyads. Letters indicate significance for the dyad at that time duration: lion-lion vs lion-hyena (a), hyena-hyena vs lion-hyena (b), lion-lion vs hyena-hyena (c), hyena-hyena vs lion-lion (d), lion-hyena vs hyena-hyena (e). Number of dyads: 9 lion-lion dyads, 4 hyena-hyena dyads, and 10 lion-hyena dyads. Statistical results are presented in S7 Table (Supporting information).

#### Movement and activity within core use areas

The activity recorded from the accelerometers of Chobe lions and spotted hyenas were significantly higher when inside the core use area of the competitor species (Fig 9), although the activity of Etosha lions and hyenas also increased when inside competitor core areas. Chobe lions had higher activity inside competitor core areas during the dusk/dawn period (mean ± SE: inside 34.29 ± 29.38 AMVs, outside 30.66 ± 25.08 AMVs; *F* = 51.31, *p* < 0.05; S8 Table), while a hyena from Chobe had higher activity both inside the competitor, and outside the conspecific core areas (competitor mean ± SE: inside 84.32 ± 1.59 AMVs, outside 65.34 ± 0.68 AMVs; conspecific mean ± SE: inside 50.97 ± 0.90 AMVs, outside 77.58 ± 0.81 AMVs; *F* = 111.7 and 433.1, respectively, all *p*-values < 0.0001; S8 Table).

**Fig 9.**
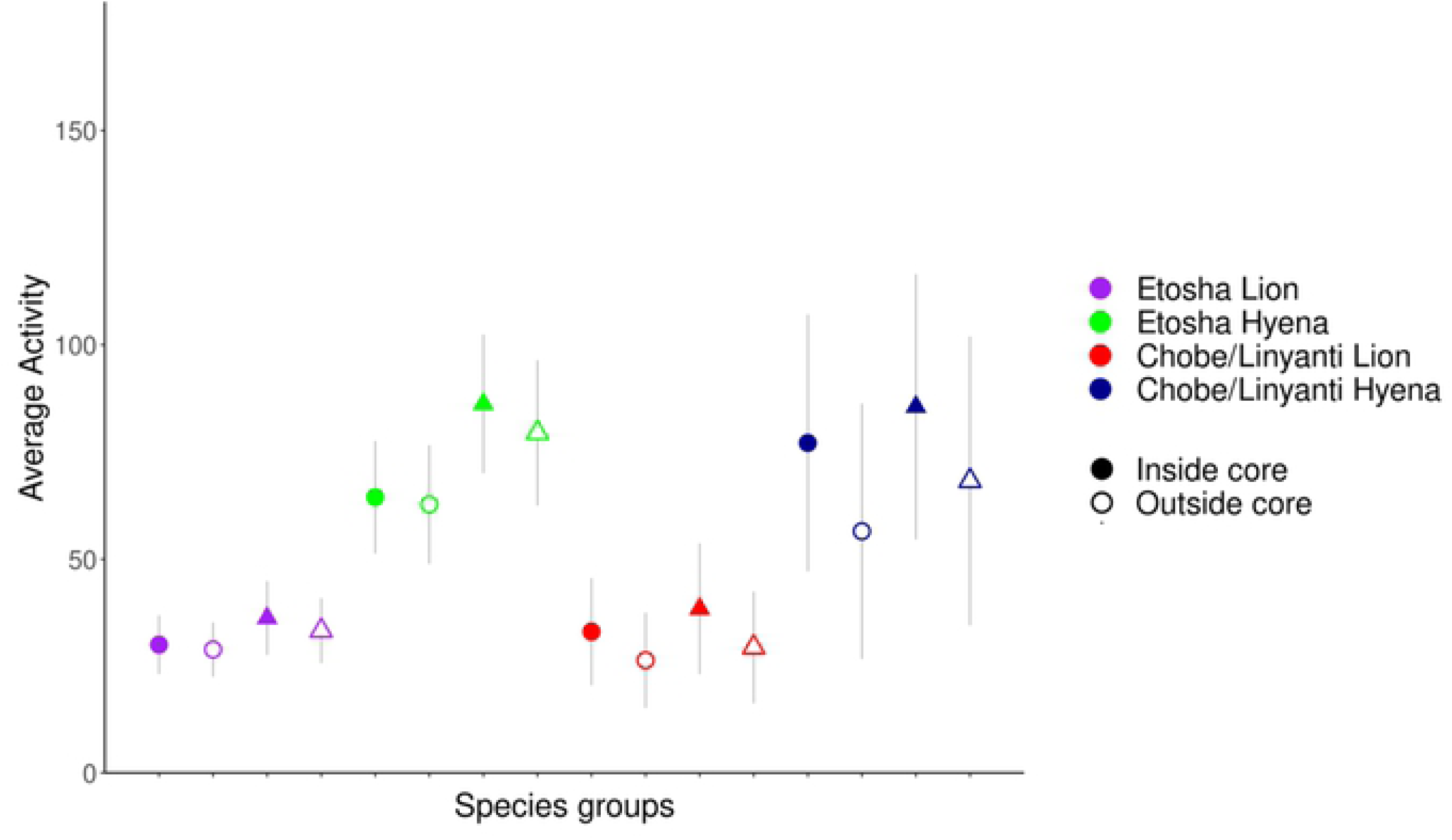
Average activity of lions and spotted hyenas with respect to competitor core use areas. Inside competitor core areas are represented by solid shapes, and outside competitor core areas are represented by open shapes. Shapes represent the mean activity and error bars the SE. Activity was measured simultaneously on each axis as the difference in acceleration between two consecutive measurements and given a relative range between 0 and 255 (activity monitor values [AMVs]), characterizing the mean activity/acceleration. Activity X (circles) = forward/backward motion. Activity Y (triangles) = rotary/sideways motion.

Additionally, lions in both ecosystems had increased activity when at closer distances to conspecifics than to competitors (Etosha mean ± SE: 100-200 m to conspecifics 56.86 ± 20.30 AMVs, to competitors 25.70 ± 20.50 AMVs, *t* = -3.77, df = 8.9, *p* < 0.01; Chobe mean ± SE: 500-600 m to conspecifics 45.39 ± 36.94 AMVs, to competitors 29.50 ± 0.01 AMVs, *t* = -80.71, df = 1, *p* < 0.01; S8 Table), with Etosha lions travelling faster (calculated from GPS locations) at closer distances to conspecifics than to competitors during dusk/dawn periods (mean ± SE: 100-200 m to conspecifics 0.412 ± 0.14 m/s, to competitors 0.082 ± 0.10 m/s, *t* = -6.78, df = 6.7; 200-300 m to conspecifics 0.337 ± 0.14 m/s, to competitors 0.155 ± 0.12 m/s, *t* = -3.78, df = 6.7; 300-400 m to conspecifics 0.302 ± 0.16 m/s, to competitors 0.085 ± 0.08 m/s, *t* = -4.96, df = 5.9; 400-500 m to conspecifics 0.396 ± 0.15 m/s, to competitors 0.135 ± 0.10 m/s, *t* = -3.25, df = 10.4; all *p*-values < 0.01; S8 Table). Similarly, hyenas from Etosha demonstrated increased activity at closer distances to conspecifics than to competitors during nocturnal periods (mean ± SE: 100-200 m to conspecifics 127.18 ± 22.53 AMVs, to competitors 47.33 ± 25.13 AMVs, *t* = 2.74, df = 5.0, *p* < 0.05), whereas they travelled at faster speeds when closer to competitors than to conspecifics during dusk/dawn periods (mean ± SE: 0-100 m to competitors 0.135 ± 0.06 m/s, to conspecifics 0.010 ± 0.004 m/s, *t* = 2.77, df = 5.1, *p* < 0.05; S8 Table). Contrarily, the dusk/dawn activity of the Chobe hyena was significantly higher at closer distances to competitors than to conspecifics (mean ± SE: 200-300 m to competitors 92.50 ± 11.59 AMVs, to conspecifics 53.06 ± 7.32 AMVs, *t* = 2.88, df = 3.9; 600-700 m to competitors 106.17 ± 11.28 AMVs, to conspecifics 59.17 ± 10.30 AMVs, *t* = 3.08, df = 15.0; all *p*-values < 0.05; S8 Table).

During dusk/dawn periods, spotted hyenas from both Etosha and Chobe travelled at faster speeds inside of competitor core areas relative to outside of competitor core areas (Etosha mean ± SE: inside 0.470 ± 0.14 m/s, outside 0.422 ± 0.14 m/s, F = 12.96, *p* < 0.05; Chobe mean ± SE: inside 0.527 ± 0.22 m/s, outside 0.363 ± 0.21 m/s, F = 8.33, *p* < 0.05; S8 Table). Similarly, hyenas from Chobe also travelled at faster speeds inside of competitor core areas during nocturnal periods (mean ± SE: inside 0.438 ± 0.13 m/s, outside 0.297 ± 0.13 m/s, F = 18.93, *p* < 0.05; S8 Table). Conversely, Chobe hyenas travelled at slower speeds when inside of conspecific core areas relative to outside of conspecific core areas during nocturnal periods (mean ± SE: inside 0.233 ± 0.23 m/s, outside 0.334 ± 0.17 m/s, *F* = 1515.36, *p* < 0.05; S8 Table). Additionally, Chobe hyenas moved at faster speeds inside competitor core areas in comparison to when they were inside conspecific core areas during nocturnal and dusk/dawn periods (nocturnal mean ± SE: competitor 0.425 ± 0.16 m/s, conspecific 0.233 ± 0.23 m/s, *t* = 6.04, df = 4.2; dusk/dawn mean ± SE: competitor 0.527 ± 0.22 m/s, conspecific 0.268 ± 0.29 m/s, *t* = 4.88, df = 4.9, all *p*-values < 0.01; S8 Table).

The tortuosity of lion movements differed when they were inside of competitor core areas in relation to when they were outside of it (Fig 10). During nocturnal periods, lions from Chobe had more directional movements inside competitor core areas (mean ± SE: inside 0.032 ± 1.08, outside -0.222 ± 0.99, Watson’s test statistic: 0.242, *p* < 0.05; S8 Table), and were also more directional inside competitor core areas than they were inside of conspecific core areas (mean ± SE: 0.113 ± 1.71; Watson’s test statistic: 0.153, 0.05 < *p* < 0.10; S8 Table). In addition, Chobe lions were less tortuous inside competitor core areas than Etosha lions were (mean ± SE: Etosha 0.055 ± 0.64, Chobe 0.032 ± 1.08, Watson’s test statistic 0.204, *p* < 0.05; S8 Table). Contrarily, Etosha lions had more tortuous movements inside competitor core areas than outside of them during dusk/dawn periods (mean ± SE: inside 0.073 ± 0.79, outside 0.029 ± 0.79, Watson’s test statistic: 0.171, 0.05 < *p* < 0.10; S8 Table). Chobe lions were also more directional inside of competitor core areas (0.032 ± 2.41) than they were inside of conspecific core areas (0.113 ± 2.88; Watson’s test statistic: 0.153, 0.05 < *p* < 0.10; S8 Table).

**Fig 10.**
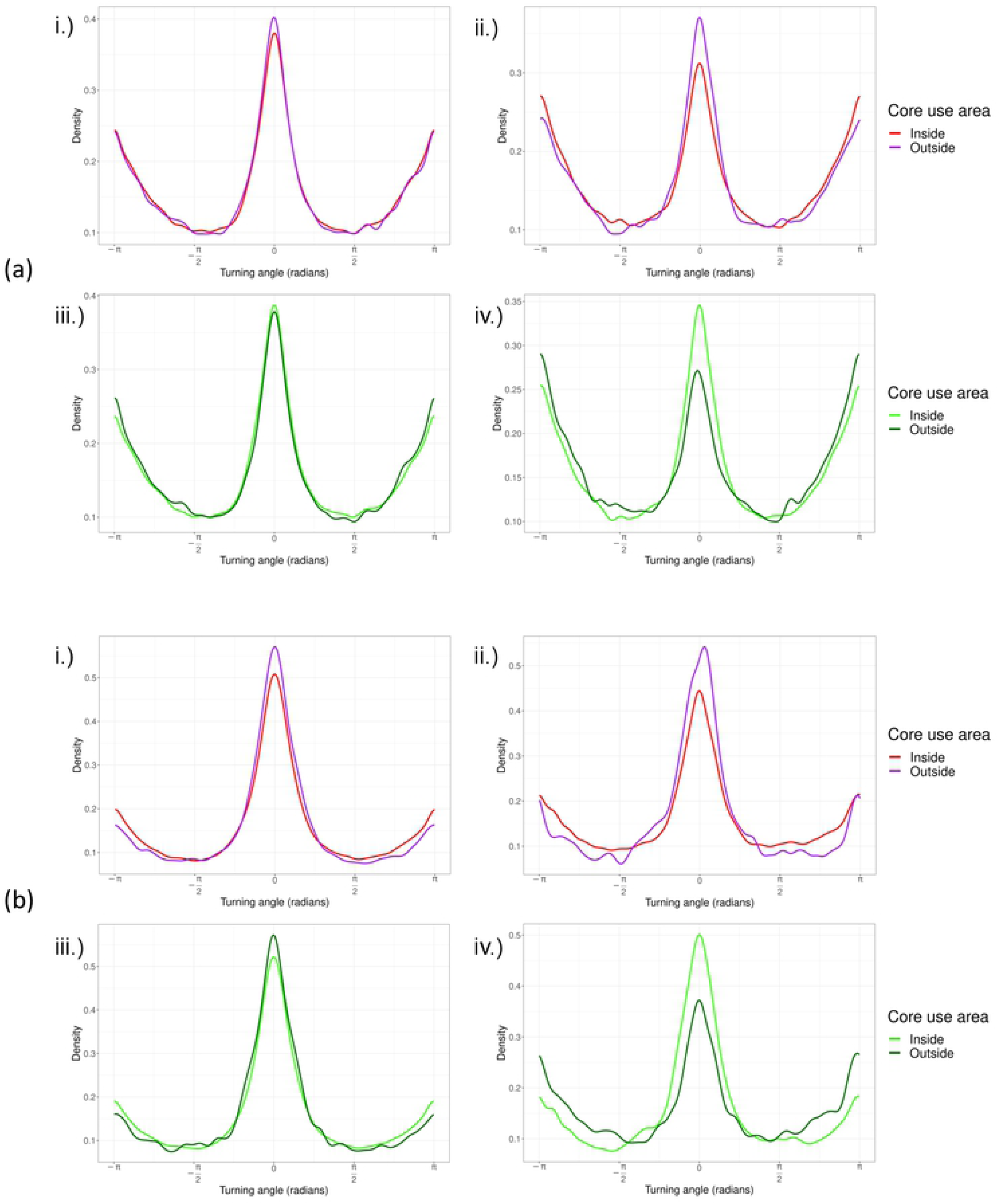
Tortuosity of lions and spotted hyenas with respect to competitor core use areas. Tortuosity of (a) lions and (b) spotted hyenas from the Etosha National Park, Namibia (i & iii) and the Chobe National Park and Linyanti Conservancy, Botswana (ii & iv). Path tortuosity is shown from inside and outside of competitor core areas (i & ii), and conspecific core areas (iii & iv).

In addition, lions from Chobe demonstrated sex-specific differences in path tortuosity with regards to conspecific core areas during the nocturnal period. Although Chobe male lions had more tortuous paths than female lions (mean ± SE: males -0.512 ± 1.74, females 0.078 ± 0.80; *F* = 7.81, *p* < 0.05), both sexes demonstrated less tortuous paths inside conspecific core areas than they did outside of them (*F* = 10.98, *p* < 0.05; S8 Table). Conversely, spotted hyenas from Etosha had more tortuous paths inside conspecific core areas with more directional movements outside conspecific core areas during nocturnal periods (mean ± SE: inside -0.079 ± 0.65, outside 0.018 ± 0.62, F = 8.65, *p* < 0.05; Fig 10; S8 Table). Furthermore, lions from Etosha had more tortuous movements when at close distances to competitors (up to 500 m) during nocturnal and dusk/dawn periods, relative to when at close distances to conspecifics (nocturnal mean ± SE: 0-100 m to competitors -2.912 ± 0.74, to conspecifics 0.169 ± 0.85, Watson’s test statistic 0.242; 100-200 m to competitors 2.775 ± 0.60, to conspecifics 0.170 ±, Watson’s test statistic 0.227; 400-500 m to competitors 2.305 ± 0.48, to conspecifics -0.001 ± 0.50, Watson’s test statistic 0.227; dusk/dawn: 100-200 m to competitors 3.130 ± 0.71, to conspecifics 0.076 ± 0.46, Watson’s test statistic 0.264; 400-500 m to competitors -2.666 ± 0.71, to conspecifics 0.058 ± 0.60, Watson’s test statistic 0.191; all *p*-values < 0.05; S3 Fig 12; S8 Table).

### Time-use metrics

Revisitation and duration (RD) space plots that indicate areas of high revisitation rates and locations of long visit durations for a lion and spotted hyena are presented in Fig 2 (with all other individuals in S4 Fig, Supporting information). The distributions of the selected variables for each of the clusters are presented alongside a map of the individual’s relocations, colour-coded according to the identified clusters within the RD space in Fig 3 (with all other individuals in S5 Fig, Supporting information). For all individuals, the factor analysis of mixed data (FAMD) method consistently selected land cover categories and time of day as high-scoring variables among the principal dimensions explaining the patterns of recursions and extended stays for each lion and spotted hyena individual. The next most consistently selected ecogeographical variables as important factors explaining the patterns of recursions and extended stays for lions were variables related to interspecific and intraspecific interactions (probability of, distance to, and whether inside of core), chosen 75% and 68.8% of the time, respectively (Table 3). However for hyenas, the most consistently selected factors among the principal dimensions were variables related to interspecific interactions (probability of, distance to, and whether inside of core), chosen 61.5% of the time. Other variables selected with equal consistency as variables of interspecific interactions for Etosha hyenas included probability of site-attracted foragers and moon illumination, while distance to permanent water was selected with equal frequency for Chobe hyenas (Table 3).

**Table 3.**
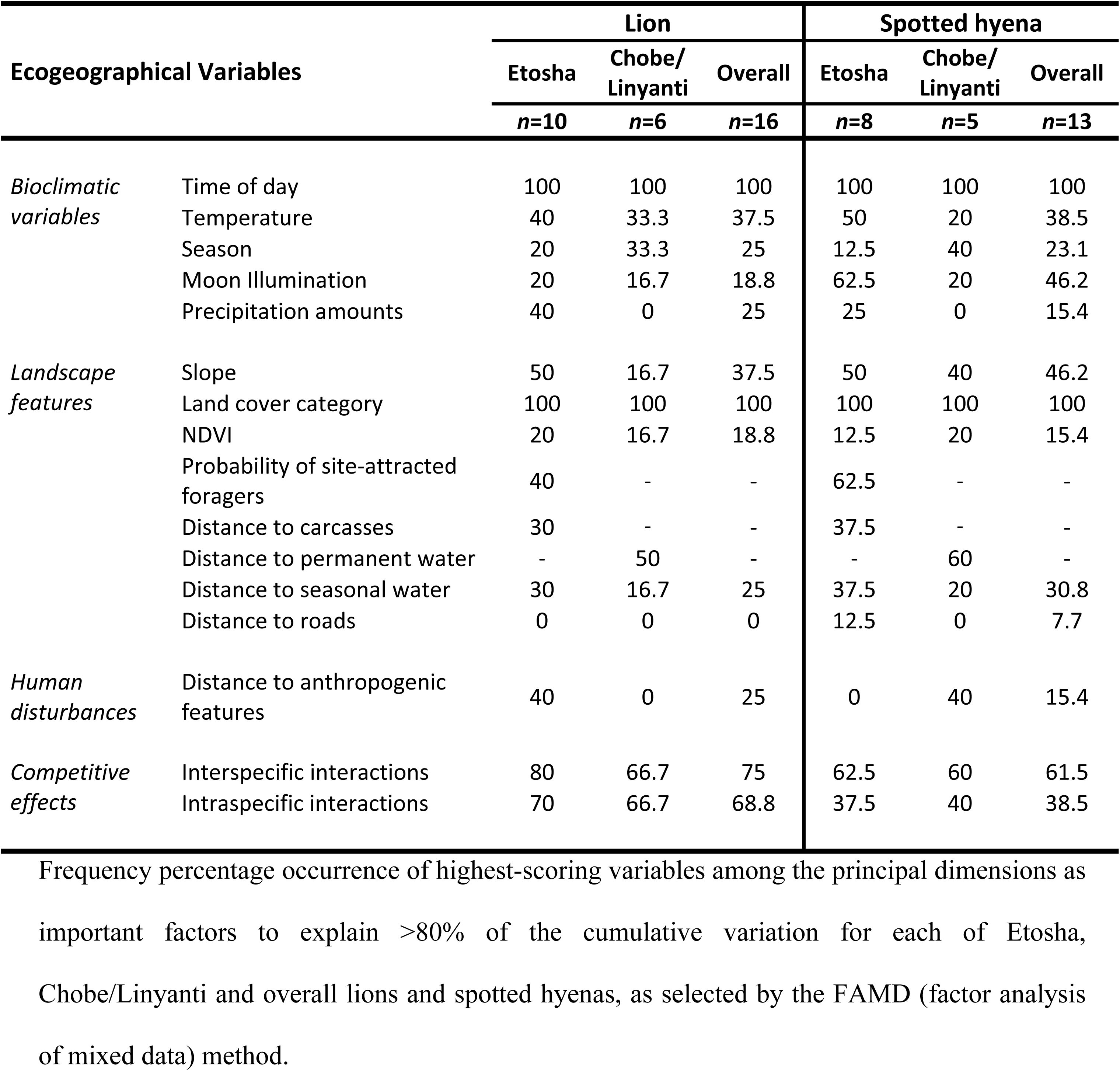
Highest-scoring ecogeographical variables for lions and spotted hyenas.

Focusing our time-use observations on variables associated with interspecific competition, we observed that lions tended to demonstrate longer durations in locations of low competitor probabilities, at far distances to competitors, and when outside competitor core use areas (see a.3 and a.8 in S5A Fig). Lions also had increased recursions in areas of high competitor probabilities, at close distances to competitors and inside competitor core use areas (all chi-square *p*-values < 0.001; Table 4). However, male lions and females of mating pairs exhibited longer durations in localities of high competitor probabilities; while single female lions had shorter durations with increased revisitations (see a.5 and c.3 versus a.4 and b.2 in S5A Fig). Similarly, spotted hyenas also tended to have longer durations in locations of low competitor probabilities, and they had higher recursions with short durations in locales of high competitor probabilities (see a.1 and b.3 in S5B Fig). Furthermore, hyenas exhibited longer durations at further distances from competitors with shorter durations and increased revisitations at shorter distances to competitors (all chi-square *p*-values < 0.001; Table 4; see a.6 and c.2 in S5B Fig).

**Table 4.**
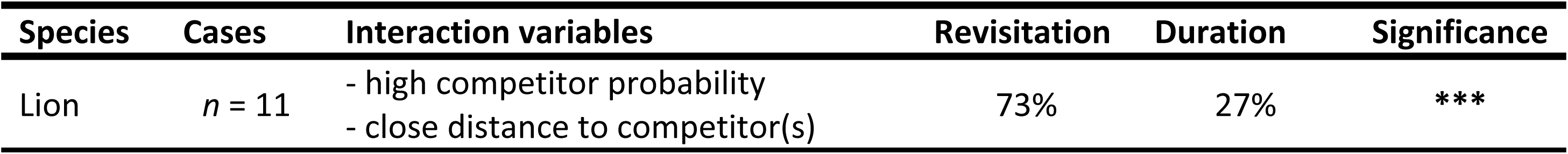
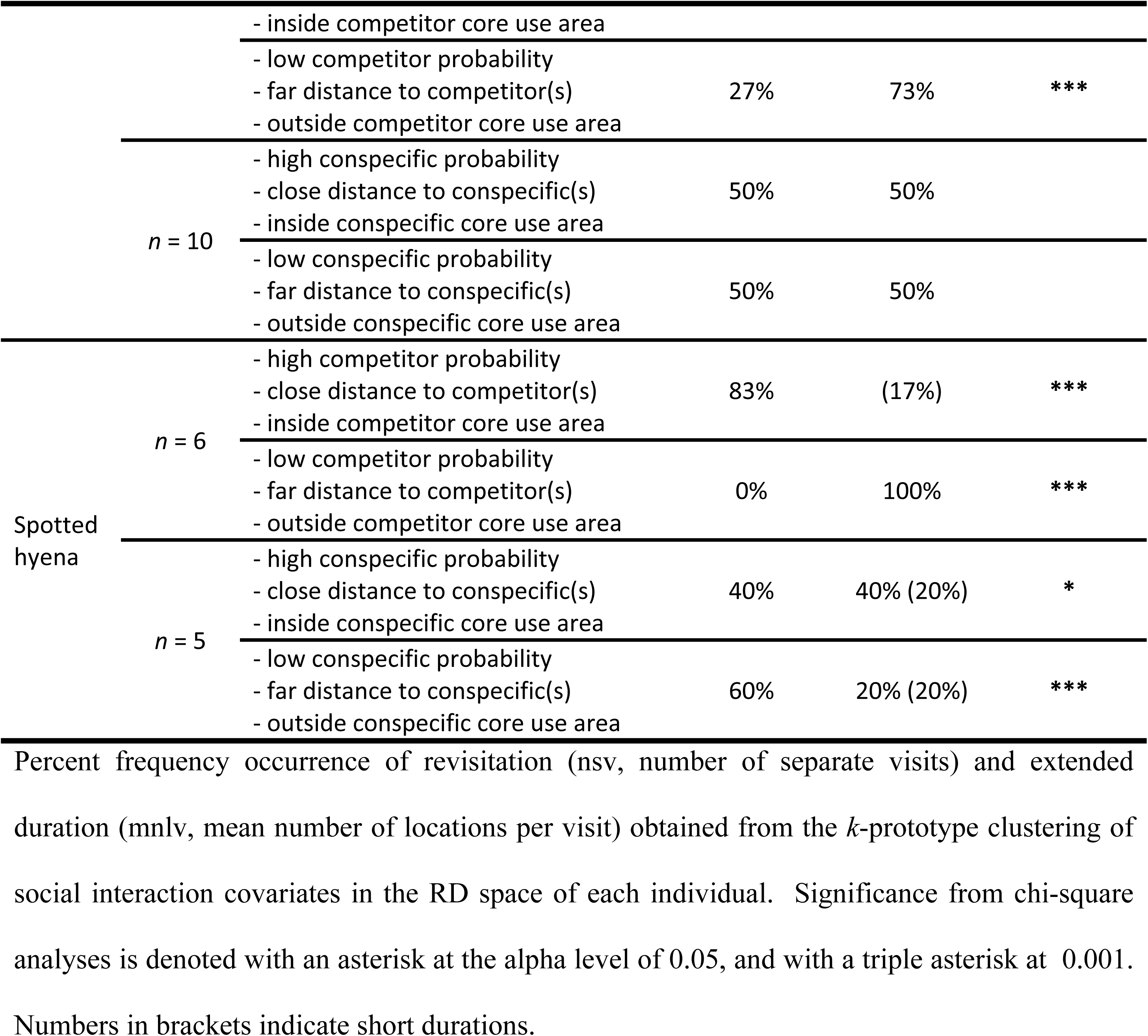
Frequency occurrence of revisitation and duration for lions and spotted hyenas according to social interaction covariates.

However, with regards to intraspecific competition, our observations revealed no differences in the time-use duration and revisitation of lions in locations of low and high conspecific probabilities (Table 4). Conversely, hyenas had twice the frequency of extended durations than they did shorter durations in locations of higher conspecific probabilities (chi-square *p*-value < 0.05; Table 4; see a.2 and a.3 in S5B Fig). In addition, hyenas also had higher recursions in localities of lower conspecific probabilities, at far distance to conspecifics, and outside conspecific core use areas (chi-square *p*-value < 0.001; Table 4).

## Discussion

Combining patterns of space use, temporal activity, fine-scale habitat use differentiation, and localized reactive avoidance behaviours in response to the potential risk of competition, has revealed the complex dynamics among lions and spotted hyenas within an apex predator system. Our findings are consistent with other studies in which lions and hyenas are often positively associated with one another [8,30,41,75], and they appear to behaviourally mediate the potential for competition by active avoidance or fine-scale behavioural mechanisms, as exhibited by other carnivores in response to the direct risk of encountering other predators [29, 92].

Environmental spatial complexity, which allows for the selection of different habitats, may promote coexistence between species [93], and has been recorded for several sympatric carnivores [14,23,36,94–100], including lions and spotted hyenas [29, 101]. However, fine-scale habitat use differentiation likely occurs as the key mechanism allowing for the coexistence of species within homogeneous landscapes [99, 102]. Therefore, while the heterogeneous landscapes of the Chobe riverfront may facilitate coexistence of species of the same trophic level [103], subtle patterns of habitat use partitioning reflected in the movement decisions of lions and spotted hyenas may further explain the persistence of the two predators across arid and mesic extremes of their environment. Our results indicate differences between lion and spotted hyena movements, which can be attributed to differences in the hunting strategies of the two species [104], potentially resulting in fine scale habitat separation rather than complete segregation [105].

Mainly cursorial predators, spotted hyenas are generally more active, travel faster and further than lions [37, 53]. As sit-and-wait predators, lions typically have lower activity, move at slower speeds, and cover shorter distances than spotted hyenas [39, 59]. Corresponding to the findings of Durant et al. [77], lions and spotted hyenas in this study utilized similar habitats, both mainly occurring within grassland habitats in Etosha and shrubland habitats in Botswana [106–108]. Despite utilizing similar habitats, how the two species exploit these habitats differs, and reflects the differences in habitat characteristics between ambush and cursorial predators [109, 110]. Lions typically use areas in which stalking cover is available [58, 111], whereas hyenas prefer to use open areas, and are generally more dispersed across the landscape [112, 113]. Likewise, Bender et al. [15] reported fine-scale habitat segregation among pumas (*Puma concolor*), coyotes (*Canis latrans*) and bobcats (*Lynx rufus*) in the San Andres Mountains as a result of preferences for habitat characteristics that facilitate movements, despite being positively associated with one another. Thus, our results lends support to a growing body of evidence that demonstrates coexistence among carnivores is facilitated by behavioural mechanisms, in addition to spatial and temporal partitioning [15,31,51,114,115].

Although the movement decisions and behavioural responses of lions and spotted hyenas are adaptable across different systems, we found that lions and spotted hyenas do not respond to inter- and intraspecific interactions equally among heterogeneous and homogeneous environments. Hyena clans in Etosha were observed to participate in more territorial clashes than they did in Chobe (unpublished data). As heterogeneous environments, or habitats of increased complexity, permit coexistence among carnivore species [5, 116], presumably the potential for competition (both inter- and intraspecific) is mitigated in Chobe [14], an environment of increased complexity [117].

Previous studies have demonstrated lion movements to be inextricably tied to the location of water-holes across the landscape due to concentrated search efforts for prey [25, 118]. In Etosha during the dry season, water is primarily supplied at developed locations from a system of pumped boreholes that are routinely serviced and maintained by park personnel. Conversely, perennial water from the Chobe river and the Okavango Delta provides a permanent water source at the Botswana sites [119]. In this study, lion core use areas either encompassed, or were anchored, by such water sources, while only half or fewer of hyena core use areas were. Our results indicate that hyenas in Chobe spent more time in areas that were closer to permanent water and had increased recursion to locations further from water. Spotted hyenas have been found to use locations far from water for den sites and resting [30, 114], thus we suggest that hyenas are potentially choosing to remain in areas with landscape characteristics that minimizes detection [120] while increasing prey vulnerability [121]. Although spotted hyenas require access to drinking water, they can survive on very little of it [38]. Presumably, hyenas in Chobe spend more time in riverine habitats which consist of relatively dense vegetative cover and greater topographic heterogeneity to increase the potential of obtaining food resources when prey aggregate at water sources [60, 122]. In areas where lions are the dominant predator, spotted hyenas may be relegated to suboptimal areas away from readily-available and prime resources, similar to coyotes and wolves (*Canis lupus*) [123]. In this way, spotted hyenas are potentially equivalent to naïve or subordinate lions that have been forced into peripheral habitats [124]. Thus, the behavioural plasticity and opportunistic behaviour in foraging and habitat use of spotted hyenas may facilitate coexistence with lions in areas where they overlap.

Interestingly, hyenas in Etosha had higher recursions to locations with higher probabilities of encountering site-attracted foragers, thus increasing their chances of coming across foraging ungulates or potentially new carcasses [125]. However, hyenas that shifted their ranges in the wet season to include the anthrax endemic areas had longer durations at these locations; presumably because they had to travel further (±60km) to access these localities. The ability to leave their home range and traverse across territories of other clans to benefit from an abundance of resources (i.e. surplus carcasses from anthrax outbreaks), is reminiscent of the commuting behaviour exhibited by spotted hyenas as they follow to exploit migratory prey during the wet season in the Serengeti [55, 126]. We suggest this behavioural plasticity is an ecological strategy that spotted hyenas appear to exploit as necessary to increase the potential appropriation of resources, thus conferring a fitness advantage.

Furthermore, recursion and duration were synonymous with travelling movements for Etosha hyenas during new moon and full moon periods. Hyena movements were more localized at higher durations during the periods between new and full moons as they spent more time foraging or searching for prey. Similar to wolves, which were documented to be nearly twice as successful when hunting on moonlit nights [127], hyenas likely require sufficient visibility to increase their hunting success [13]. During this study, hyenas were observed to undertake cursorial hunts during moonlit nights, and switched to an ambush strategy coupled with opportunistic chases during darker periods. Comparably, lions typically experience higher hunting success during dark nights, although they were less successful at appropriating prey during moonlit nights [128]. In addition, hyenas were observed (during night follows in this study) to rest when the moon was at its brightest during full moon nights, and would only resume foraging activities after the moon had lowered in the sky (pers. obs.). This suggests that hyenas are likely to focus on directed movements (traversing between patches) during dark periods, and resting when it is too bright to avoid detection by prey [129]. Accordingly, hyenas focus their foraging and hunting efforts during periods of sufficient light conditions between the time of new and full moon nights, allowing for avoidance of the potential risk of interspecific encounters with lions [130, 131].

Temporal heterogeneity in conjunction with spatial heterogeneity likely occurs as a mitigation strategy in response to interference competition, and facilitates coexistence among carnivores [36, 132]. Our findings indicate lions and spotted hyenas were both nocturnal, with the lion more diurnal than the hyena. Similarly, Sogbohossou et al. [101] found the activity of lions and spotted hyenas to be spread over the night with no real peaks. However, we found evidence of temporal partitioning on a finer scale than nocturnal and crepuscular patterns, as has been recorded in other studies [13,26,31,133]. Thus, our results correspond with studies in which temporal partitioning occurred as a result of differences in activity periods between predator species, and was presumed to be the main driver for coexistence in sympatric carnivores [3,9,95].

### Intraspecific and interspecific interactions

Since lions tend to exhibit a high degree of coordinated movements among lion pairs [134], intraspecific interactions were also selected as important factors accounting for the time-use patterns among lions. However, as we did not differentiate between competitive or mutually beneficial intraspecific interactions, it is plausible that mating pairs may have exaggerated or confounded this result. However, interspecific interactions were important influences on the time-use metrics that drives both lion and spotted hyena space-use patterns. Surprisingly, although female lions had shorter durations in locations with high competitor probabilities, we found that male lions (and their mating partners) had longer durations in these areas. As dominant predators, male lions are either unaffected by being in locations of higher competitor probabilities, or it is likely that they spend more time within these areas to increase the likelihood of encountering hyenas, because they appear to derive benefits from hyenas through kleptoparasitism [43,135–138]. During this study, male lions were often observed to be attracted to spotted hyena calls: they would immediately perk up their ears, turn to face the direction of the calls, and often walked towards the direction of the hyena sounds. Observations of direct interactions between lions and spotted hyenas during this study almost always involved hyenas actively avoiding male lions by retreating and moving away, as was also observed in other studies [43, 135]. At encounters with male lions at fresh kills, hyenas often lost their kills to lions or aggregated into large groups (>20 individuals) before attempting to initiate mobbing behaviour, similar to the findings of previous studies [106, 139].

In addition, our results demonstrated patterns of recursions and locations of extended stays within the home ranges of apex predators to be influenced by the probability of, and proximity to, competitor and conspecifics. We suggest this to be the result of a perceived risk of competition, in which carnivores behaviourally mediate the potential for competition by altering their space use, or movement patterns [27]. Furthermore, other studies have documented the behavioural response of carnivore species to the potential risk of either encountering competitors or competitive interactions with increased vigilance and movements, either in preparation for potential interactions, or to move through/exit the area as quickly as possible [29,50,140]. Pangle and Holekamp [141] attributed the vigilance levels of spotted hyenas to be more influenced by interspecific than intraspecific threats. Likewise, Valeix et al. [142] documented lions moving at quicker speeds, and with relatively straighter trajectories, in response to the risk of conflict when close to human settlements. Specifically, we found spotted hyenas to behaviourally reduce the risk of potential interaction with lions by remaining longer in areas of low competitor probabilities and at far distances to competitors. This type of behaviour is mirrored in the behaviour-specific habitat selection of lions’ in response to mitigating the potential risk of conflict with humans [143]. Thus, the behavioural responses of lions and spotted hyenas towards the perceived risk of competition appears to be similar, regardless of whether it stems from inter- or intraspecific interactions.

## Conclusions

Our results have implications for the conservation of large carnivores in substantiating the potential effects of interference competition on lion and spotted hyena spatial patterns and movements. These findings supplement the growing body of evidence that demonstrates coexistence among carnivores is facilitated by fine-scale behavioural mechanisms in addition to spatial and temporal partitioning. While the patterns of spatial and temporal overlap observed among lions and hyenas do not differ from earlier studies; combining patterns of space use, temporal activity, fine-scale habitat use differentiation, and localized reactive avoidance behaviours in response to the potential risk of competition, has revealed the complex dynamics among lions and spotted hyenas within an apex predator system. Additionally, patterns of recursion and locations of extended stays within the home ranges of apex predators are influenced by the probability of and proximity to competitors and conspecifics, and can be used to inform management strategies for the maintenance of carnivore communities.

As large carnivores are becoming increasingly constrained to protected areas, it is important to note that lions and spotted hyenas do not respond to inter- and intraspecific interactions equally among heterogeneous and homogeneous environments. Specifically, the movement decisions and behavioural responses of lions and spotted hyenas are adaptable across different systems, and are likely a result of several factors, including habitat complexity, hunting strategies, and the active avoidance of interspecific and heterospecific competitors. Consequently, we encourage conservation practitioners to recognize the importance of the potential effects of inter- and intraspecific interactions among apex predators in managing diverse, ecological communities.

## Acknowledgements

We thank Dr. Frans Joubert, Christopher Diab, Danica Metlay, Jacqueline Moser, Julie Fabricius Faustrup, Astri Frafjord, Linda Molloy, Christopher Mastropietro for dedicated field work, and Marthin Kasaona, Gabriel Shatumbu, Shayne Kotting, Mathews Daniel, Fredrick Khumub, Malakia Hango, Isaskar Uahoo, Dr. Rob Jackson, Dr. Caron Botes, and Dr. Larry Patterson for support and assistance in the field. We also thank the Ministry of Environment and Tourism Namibia and Boas Erckie (Etosha National Park), the Department of Wildlife and National Parks Botswana and Dr. Michael Flyman (Chobe National Park), the Chobe Enclave Community Trust (Linyanti Conservancy), the Okavango Kopano Mokoro Community Trust and S.K. Moepedi (OKMCT), Sanctuary Retreats and Charl Badenhorst for infrastructure support and permissions. We are grateful to the Etosha Ecological Institute and CARACAL Biodiversity Center for the use of lab facilities, and to Dr. Richard Fynn and the Okavango Research Institute for materials. We thank Chris Coetzee and Fritz Stolzenberg in Namibia, and Trish Williams in Botswana for support outside the parks. We are indebted to Drs. Jerry and Jana Lackey for their constant and unwavering support. We are also grateful to Lotek Wireless Inc., Shush Productions, African Lion and Environmental Research Trust (ALERT), the Namibian Environment & Wildlife Society (NEWS), Gert Uls and Resun GmbH for sponsoring some of the field equipment. We thank Helicopter Horizons for their assistance with animal captures, and to Dr. John ‘Tico’ McNutt and Mike Holding for their assistance with flight-tracking. We also thank Andy Lyons for assistance with T-LoCoH, Dana Seidel for guidance with coding R analyses, and Dr. Wolfgang Beyer for undertaking serological work.

## Supporting information

**S1 Appendix. Additional details on study sites and data collection.**

**S2 Appendix. Descriptive analyses of lion and spotted hyena movement descriptors.**

**S1 Table. Percent frequency of lion and spotted hyena relocations by land cover type.** Relocations were recorded from lions and spotted hyenas in the Etosha National Park, Namibia, the Chobe National Park and Linyanti Conservancy, Botswana, and only from lions in the NG32 concession of the Okavango Delta, Botswana.

**S2 Table. Seasonal utilization distributions (UDs) for all collared lion and spotted hyena individuals.** Kernel density (a, c) and *a*-LoCoH (b, d) area measures for home ranges and core use areas (km^2^) of lion (a, b) and spotted hyena (c, d) individuals. Upper panels consists of lion and hyena individuals from the Etosha National Park, Namibia, with lower panels from the Chobe National Park and Linyanti Conservancy. For lions, the NG32 concession in the Okavango Delta, Botswana in also included. CS = combined seasons, DS = dry season, WS = wet season.

**S3 Table. Areas of overlap in lion and spotted hyena ranges.** Total overlapped areas in km^2^ between lions’ (vertical column) and spotted hyenas’ (horizontal column) home ranges and core areas in the (a) Etosha National Park, Namibia; (b) Chobe National Park and Linyanti Conservancy, Botswana. Utilization distributions were generated with the home range (95%) and core use area (50%) kernel density estimator (i) and *a*-LoCoH (ii) isopleths. Males are underlined. An asterisk denotes mortality.

**S4A Table. Proportion of overlaps with spotted hyenas in lion ranges.** Total proportion of lion (vertical column) home ranges and core areas overlapped by spotted hyena individuals (horizontal column) in the (a) Etosha National Park, Namibia; (b) Chobe National Park and Linyanti Conservancy, Botswana. Utilization distributions were generated with the home range (95%) and core use area (50%) kernel density estimator (i) and *a*-LoCoH (ii) isopleths. Males are underlined. An asterisk denotes mortality.

**S4B Table. Proportion of overlaps with lions in spotted hyena ranges.** Total proportion of spotted hyena (vertical column) home ranges and core areas overlapped by lion individuals (horizontal column) in the (a) Etosha National Park, Namibia; (b) Chobe National Park and Linyanti Conservancy, Botswana. Utilization distributions were generated with the home range (95%) and core use area (50%) kernel density estimator (i) and *a*-LoCoH (ii) isopleths. Males are underlined. An asterisk denotes mortality.

**S5 Table 1. Lion and spotted hyena movement metrics.** (a) all lions and spotted hyenas, and (b) lions and spotted hyenas from the Etosha National Park, Namibia (ENP) and the Chobe National Park, Linyanti Conservancy, and the NG32 concession of the Okavango Delta^†^, Botswana (CNP). Movement metrics consist of the means ± standard deviations for activity (activity monitor values [AMVs]), step length (m), speed (m/s), net-squared displacement (NSD, km^2^), and path tortuosity (radian).

^†^*No spotted hyenas were collared from the Okavango Delta, Botswana*.

**S5 Table 2. Lion and spotted hyena activity and movement metrics in relation to the lunar cycle.** Activity (AMVs), step length (m), and path tortuosity (radian) of lions and spotted hyenas from the Etosha National Park, Namibia (ENP), and the Chobe National Park, Linyanti Conservancy, and the NG32 concession of the Okavango Delta^†^, Botswana (CNP). Values are means ± standard deviations during the nocturnal (30min fixes from 18h00-6h00 and 17h00-8h00) and dusk/dawn (5min fixes from 19h00-21h00 and 4h00-6h00) periods.

^†^*No spotted hyenas were collared from the Okavango Delta, Botswana*.

**S5 Table 3. New and full moon activity of lions and spotted hyenas from the 24-hour period cycle.** Activity (AMVs) of lions and spotted hyenas from the Etosha National Park, Namibia (ENP), and the Chobe National Park, Linyanti Conservancy, and the NG32 concession of the Okavango Delta^†^, Botswana (CNP). The 24-hour cycle was subdivided into seven different periods. Night/nadir/night end consists of the end of evening twilight to the beginning of morning twilight divided into three equal intervals. Afternoon = noon to sundown; dusk = sundown to twilight end; dawn = beginning of morning twilight to sunrise; morning = sunrise to noon. Values indicate the means ± standard deviations of the seven different periods of the 24-hour cycle during new and full moon phases of the dry and wet seasons.

^†^*No spotted hyenas were collared from the Okavango Delta, Botswana*.

**S5 Table 4. Lion and spotted hyena movement metrics according to low, medium, and high body condition scores.** Speed (m/s) and path tortuosity (radian) of lions and spotted hyenas during nocturnal (30min fixes) and dusk/dawn (5min fixes) periods from the Etosha National Park, Namibia, the Chobe National Park, Linyanti Conservancy, and the NG32 concession of the Okavango Delta^†^, Botswana. Values are indicated in means ± standard deviations. Body conditions of individuals were scored from spinal palpations of immobilized individuals during capture events.

^†^*No spotted hyenas were collared from the Okavango Delta, Botswana*.

**S5 Table 5. Lion and spotted hyena movement metrics according to site-attracted foragers in the Etosha National Park, Namibia.** Speed (m/s) and path tortuosity (radian) of lions and spotted hyenas during nocturnal (30min fixes) and dusk/dawn (5min fixes) periods according to the probability of site-attracted foragers (i.e. ungulates) in sites that consisted of anthrax positive carcasses from previous years.

**S6 Table. Statistical results to accompany Fig 7 in the main text.** Chi-square and *t*-test results for the percent frequency occurrence of, and average distance in meters to, the nearest conspecific and competitor for each bin of distance intervals. An asterisk denotes significance at the alpha level with * < 0.05, ** < 0.01, *** < 0.005, and **** < 0.001.

**S7 Table. Statistical results to accompany Fig 8 in the main text.** *T*-tests of consecutive time points among dyads (indicating longer time duration) at various distance intervals (a), and the frequency occurrences of dyads for different consecutive time points (b). An asterisk denotes significance at the alpha level with * < 0.05, ** < 0.01, *** < 0.005, and **** < 0.001.

**S8 Table. Movement and activity metrics within competitor and conspecific core use areas.**

Nocturnal (30min fixes) and dusk/dawn (5min fixes) periods of lion and spotted hyena activity (AMVs), speed (m/s), and path tortuosity (radian) from (a) inside and outside of competitor and conspecific core use areas, and (b) at distance intervals in meters to the nearest competitor and conspecific. Collared individuals were from the Etosha National Park, Namibia (ENP), the Chobe National Park and Linyanti Conservancy, Botswana (CNP).

**S1 Fig. Temporal schedule of collar overlap for lions and spotted hyenas.** Individuals of the Etosha National Park, Namibia (left) are separated from individuals of the Chobe National Park, Linyanti Conservancy, and Okavango Delta, Botswana (right) at the bold line indicated on 6 April 2015. Males are denoted with an asterisk.

**S2A Fig. Lion nocturnal space use.** Utilization distributions were constructed with the KDE (far left and second from right) and LoCoH a-method (second from left and far right). Panels represent the 95% isopleth of the individual’s home range for the dry season (left two panels) and wet season (right two panels). Unique identifiers are depicted vertically on the left of each row of maps. Maps indicate the individual’s home range on a satellite image of the (a) Etosha National Park with the salt pan visible; (b) Chobe National Park with the Chobe river from west to east; (c) Linyanti Conservancy with the Linyanti river from southwest to northeast; and the (d) NG32 concession in the Okavango Delta on the southwestern tip of Chief’s Island. Map source: Google Imagery, TerraMetrics.

**S2B Fig. Spotted hyena nocturnal space use.** Utilization distributions were constructed with the KDE (far left and second from right) and LoCoH a-method (second from left and far right). Panels represent the 95% isopleth of the individual’s home range for the dry season (left two panels) and wet season (right two panels). Unique identifiers are depicted vertically on the left of each row of maps. Maps indicate the individual’s home range on a satellite image of the (a) Etosha National Park with the salt pan visible; (b) Chobe National Park with the Chobe river from west to east; (c) Linyanti Conservancy with the Linyanti river from southwest to northeast. Map source: Google Imagery, TerraMetrics.

**S3 Fig 1-12. Compilation of lion and spotted hyena movement descriptors.** Movement descriptors of lion and spotted hyenas. Parameters consist of (1) speed, (2-5) step lengths, (6-7) net squared displacements, and (8-12) turning angles.

**S4A Fig. Lion revisitation and duration (RD) space plots.** Time-use constructs of lion individuals from the (a) Etosha National Park, Namibia; (b) Chobe National Park; (c) Linyanti Conservancy; and (d) Okavango Delta, Botswana. Unique identifiers are depicted vertically on the left of each row of figures. α-LoCoH hulls of individual’s utilization distributions (far left). Hull parent points coloured by visitation rate (nsv, number of separate visits; second from left), and duration of visit (mnlv, mean number of locations in the hull per visit; third from left). RD space scatterplots (second from right) with X-axis = visitation rate (nsv), and Y-axis = duration of visit (mnlv), provide a legend for revisitation/duration (RD) values for the map (far right). Points in the RD space have been jiggled to better see point density, and each point represents a hull. Points on the maps are coloured by their location in the RD space. Separate visits are defined by an inter-visit gap period ≥ 12 hours. Hulls were created using the adaptive method. Duplicate points are offset by 1 map unit.

**S4B Fig. Spotted hyena revisitation and duration (RD) space plots.** Time-use constructs of spotted hyena individuals from the (a) Etosha National Park, Namibia; (b) Chobe National Park; and (c) Linyanti Conservancy, Botswana. Unique identifiers are depicted vertically on the left of each row of figures. α-LoCoH hulls of individual’s utilization distributions (far left). Hull parent points coloured by visitation rate (nsv, number of separate visits; second from left), and duration of visit (mnlv, mean number of locations in the hull per visit; third from left). RD space scatterplots (second from right) with X-axis = visitation rate (nsv), and Y-axis = duration of visit (mnlv), provide a legend for revisitation/duration (RD) values for the map (far right). Points in the RD space have been jiggled to better see point density, and each point represents a hull. Points on the maps are coloured by their location in the RD space. Separate visits are defined by an inter-visit gap period ≥ 12hours. Hulls were created using the adaptive method. Duplicate points are offset by 1 map unit.

**S5A Fig. Cluster analyses of lion revisitation and duration.** Maps (left panels) depict the individual lion’s relocations as four (top row), six (middle row), and eight (bottom row) clusters in the (a) Etosha National Park, Namibia; (b) Chobe National Park; (c) Linyanti Conservancy; and (d) Okavango Delta, Botswana. Unique identifiers are depicted on top corner of each page. Relocations are colour-coded according to the clusters indicated by the range of revisitation (number of separate visits) and duration (mean number of locations per visit) values in RD space plots (shown in central panels). Clusters in the RD space were determined with the *k*-prototype algorithm and are based on ecogeographical variables attached to each relocation. The smaller plots (right panels) present the distribution and percent category of clusters for each of the ecogeographical variables selected from the factor analysis of mixed data (FAMD).

**S5B Fig. Cluster analyses of spotted hyena revisitation and duration.** Maps (left panels) depict the individual spotted hyena’s relocations as four (top row), six (middle row), and eight (bottom row) clusters in the (a) Etosha National Park, Namibia; (b) Chobe National Park; and (c) Linyanti Conservancy, Botswana. Unique identifiers are depicted on top corner of each page. Relocations are colour-coded according to the clusters indicated by the range of revisitation (number of separate visits) and duration (mean number of locations per visit) values in RD space plots (shown in central panels). Clusters in the RD space were determined with the *k*-prototype algorithm and are based on ecogeographical variables attached to each relocation. The smaller plots (right panels) present the distribution and percent category of clusters for each of the ecogeographical variables selected from the factor analysis of mixed data (FAMD).

